# Hippocampal Synchrony Dynamically Gates Cortical Connectivity Across Brain States

**DOI:** 10.1101/2025.07.27.667035

**Authors:** G. Lazcano, M. Caneo, A. Aguilera, G. Fernandez, A. Lara-Vasquez, N. Espinosa, P. Fuentealba

## Abstract

Memory consolidation is thought to rely on hippocampo-cortical dialogue orchestrated by three cardinal sleep oscillations: cortical slow oscillations, thalamic spindles, and hippocampal sharp-wave ripples. However, how hippocampal outputs are routed to specific cortical targets and dynamically regulated across brain states remains incompletely understood. Here, we performed simultaneous multisite recordings from the dorsal and ventral hippocampus, and frontal and parietal cortex in rats alternating between wakefulness and sleep. Frontal slow oscillations operated as a global clock, resetting thalamic circuits and initiating spindle volleys that propagated from anterior to posterior cortex, while parietal slow oscillations more effectively recruited hippocampal ripples. Hippocampal ripples reflected anatomical connectivity, as dorsal ripples preferentially enhanced parietal spindles, whereas ventral ripples engaged mainly frontal spindling. Notably, when ripples synchronized in dorsal and ventral hippocampus, local excitatory drive sharply decreased and neuronal spiking redistributed, associated cortical slow oscillations and spindle responses diminished, and cortical neuronal reactivation was suppressed, indicating that dorso-ventral ripple synchrony gates, rather than amplifies, hippocampo-cortical communication. This gating effect was most evident through interactions with brain state, as dorsal-driven reactivation persisted across vigilance states, while ventral pathways were more pronounced during sleep. Collectively, our results outline a multilayer architecture in which slow oscillations provide a global temporal scaffold, spindles implement anatomically specific reactivation channels, and ripple coordination gates hippocampo-cortical communication, likely shaping the precision and specificity of memory consolidation within a highly variable neural substrate.

**Significance:** Sharp-wave ripples in the hippocampus are critical for memory consolidation, in part due to their precise temporal coordination with cortico-thalamic sleep rhythms. However, whether ripple synchronization along the hippocampal septo-temporal axis modulates cortical memory processing has remained unresolved. Here, we show that coordinated ripples, occurring simultaneously in dorsal and ventral CA1, are observed more frequently than predicted by independent occurrence and are associated with significant suppression of cortical reactivation compared to isolated episodes. This suppression does not reflect simple changes in ripple structure or increased inhibition, but instead is linked to a redistribution of excitatory drive across the hippocampal septo-temporal axis. Importantly, this gating effect is present across both sleep and wakefulness. These findings suggest that hippocampal synchrony dynamically gates, rather than amplifies, cortical engagement, thus refining systems consolidation models by highlighting the significance of network organization and precise temporal dynamics in shaping memory replay.

## INTRODUCTION

Memory consolidation critically depends on precise interactions between cortical and hippocampal networks during sleep (1), orchestrated by canonical sleep oscillations: cortical slow oscillations, thalamocortical spindles, and hippocampal sharp-wave ripples (2). These oscillations are hierarchically organized during sleep (3), and, according to the active systems consolidation model (4), this nested structure supports memory consolidation by allowing slow oscillations to temporally organize spindles, which in turn facilitate the emergence of ripples, thereby enabling effective hippocampo-cortical communication (1). Although the temporal coordination among these rhythms is well-established (2, 5), their anatomical specificity and the dynamics of their propagation across distinct brain regions remain largely unexplored.

Additionally, the hippocampus exhibits significant functional heterogeneity along its septo-temporal axis (6). Extensive research has demonstrated a clear functional division, with the dorsal CA1 (CA1d) primarily involved in spatial navigation and contextual memory encoding, whereas the ventral CA1 (CA1v) predominantly participates in emotional processing and motivational behavior (7–9). These hippocampal domains differ substantially in their intrinsic circuits and extrinsic connectivity; CA1d robustly innervates regions such as the retrosplenial cortex (RSC), posterior parietal cortex, and dorsal subiculum (10–12), while CA1v projects extensively to the medial prefrontal cortex (PFC), nucleus accumbens, and amygdala (13–15). This anatomical specialization suggests that ripple events originating from different hippocampal domains could selectively engage distinct cortical networks. Indeed, ripple oscillations can arise at various positions along the hippocampal septo-temporal axis, spreading septally or temporally, thus potentially activating different cortical circuits (16). However, it remains unknown whether this directional propagation affects cortical processing, particularly regarding memory-related neural activity patterns in cortical target areas.

Ripple events have traditionally been considered central to memory reactivation and consolidation processes (17–19). More recent findings demonstrate that hippocampal ripples can couple with neocortical ripples, thereby facilitating memory trace transfer from hippocampal to association cortical networks (20–22). In the hippocampus, ripples can occasionally occur simultaneously across the dorsal and ventral poles (7, 16, 23), although their incidence is seemingly low and the implications of their coordination remain unclear. According to the active systems consolidation framework, coordinated ripple events should enhance cortical reactivation by promoting broader hippocampal-cortical interaction (24). Contrary to these predictions, our findings indicate a larger coincidence than expected merely by chance, and a paradoxical effect of hippocampal ripple synchrony: instead of amplifying cortical reactivation, dorsal-ventral ripple coordination appears to redistribute hippocampal activity and consequently suppress cortical engagement. These unexpected observations underscore how coordinated ripple episodes dynamically modulate hippocampo-cortical interactions during sleep-mediated memory consolidation, revealing that hippocampal ripples function not as broad-spectrum facilitators, but rather as selective gating mechanisms that refine cortical memory processing.

## RESULTS

### Hierarchical organization of sleep oscillations

To characterize the interaction structure of canonical sleep oscillations, we performed simultaneous recordings of neuronal activity from anatomically interconnected hippocampal (CA1d and CA1v) and cortical regions (RSC and PFC, **Table S1**, **Fig. S1**) in rats resting after performing a spatial task that engages hippocampal-prefrontal interactions. Neural activity was monitored across interleaved task and sleep sessions (**Table S2**). Hence, we extracted cortical slow oscillations (SOs), cortical spindles, and hippocampal sharp-wave ripples to assess their precise temporal coordination (**Fig. 1a**). SOs appeared to be the master clock for hippocampo-cortical dynamics. Aligning spindle probability to the PFC SO onset (i.e.; down-state) revealed a biphasic sequence in which both PFC and RSC spindling dipped below baseline just before the onset (–1.35 ± 0.1 z) and then rebounded in a high-gain burst with similar dynamics, but with larger amplitude frontally than parietally (PFC, 5.45 ± 0.3; RSC, 4.23 ± 0.2 z; Wilcoxon signed-rank test, FDR-corrected, p < 10^-4^, **Fig. 1b**). When the reference was switched to the RSC SO, the trough was practically abolished, and the subsequent spindle burst was largely reduced, but still slightly larger frontally (PFC, 2.64 ± 0.2; RSC, 2.21 ± 0.2 z; Wilcoxon signed-rank test, FDR-corrected, p < 10^-3^, **Fig. 1b**). Overall, SO evoked stronger spindling in the frontal cortex (two-way ANOVA, F(1,521) = 172.7, p = 2.8×10^-24^; post-hoc Tukey comparison, p = 1.1×10^-10^), indicating that frontal waves broadcast globally whereas parietal waves tune activity locally. Spindle onset latencies relative to SOs were comparable between regions (two-way ANOVA, F(1,521) = 2.0, p = 0.15; post-hoc Tukey comparison, p = 0.15), and consistently positive (Wilcoxon signed-rank test against zero, z = 19.8, 343.75 ± 7.2 ms, p = 2.5×10^-87^). In addition, SOs biased ripple generation with distinct septo-temporal preferences. PFC SOs were preceded by a gradual ripple buildup culminating in a brief dorsal-larger peak at the SO onset (CA1d, 2.29 ± 0.2 z; CA1v, 0.94 ± 0.2 z, Wilcoxon signed-rank test, FDR-corrected, p < 10^-3^; **Fig. 1c**). RSC SOs recruited both hippocampal poles with similar dynamics, yet again dorsal hippocampus responses dominated (CA1d, 3.22 ± 0.2 z; CA1v, 2.47 ± 0.3 z; Wilcoxon signed-rank test, FDR-corrected, p < 0.001; **Fig. 1c**). Overall, SO-evoked ripple bursts were stronger in the dorsal when compared to ventral hippocampus (two-way ANOVA, F(1,412) = 22.4, p = 3.0×10^-6^; post-hoc Tukey comparison, p = 2.2×10^-6^), suggesting that SO preferentially propagated into the dorsal hippocampus. Ripple onset latencies relative to SOs were shorter in the dorsal compared to the ventral hippocampus (two-way ANOVA, F(1,412) = 5.3 p = 0.02; post-hoc Tukey comparison, p = 0.02), suggesting that SO reaches earlier the dorsal hippocampal pole. To further confirm this hierarchical sequence, we then used hippocampal ripples as a reference point and analyzed subsequent cortical spindle activity (**Fig. 1d**). Dorsal ripples preferentially launched parietal spindle bursts while only modestly engaging frontal spindles (RSC, 2.48 ± 0.1 z; PFC, 1.08 ± 0.1 z; Wilcoxon signed-rank test, FDR-corrected, p < 10^-3^; **Fig. 1d**), whereas ventral ripples produced cortical responses with similar time course, yet dominated by the frontal cortex (PFC, 2.35 ± 0.2 z; RSC, 1.70 ± 0.2 z, Wilcoxon signed-rank test, FDR-corrected, p < 10^-4^; **Fig. 1d**), consistent with known anatomical hippocampo-cortical connectivity patterns (25). Spindle latencies relative to ripple onsets were consistently longer in frontal than parietal cortex (PFC, 332.9 ± 57.1 ms; RSC, −20.7 ± 53.2 ms, Wilcoxon signed rank test, z = 4.9, p = 8.5×10^-7^), matching anatomical distance. In addition, sleep oscillations exhibited robust cross-frequency phase modulation. Indeed, SOs strongly modulated spindle occurrence, with spindles preferentially emerging during the up-state, and frontal spindles consistently leading parietal spindles (**Fig. S2**). Cortical spindles modestly but significantly modulated hippocampal ripple incidence, with ripples most frequently occurring around the trough of the spindle cycle (**Fig. S2**). Collectively, these findings establish that frontal SOs operate as a global temporal scaffold, parietal SOs selectively tune hippocampal engagement, and ripple–spindle coupling reflects direct anatomical connectivity, together revealing a hierarchical and topographically organized oscillatory framework that structures hippocampo-cortical communication during sleep.

**Figure 1.**
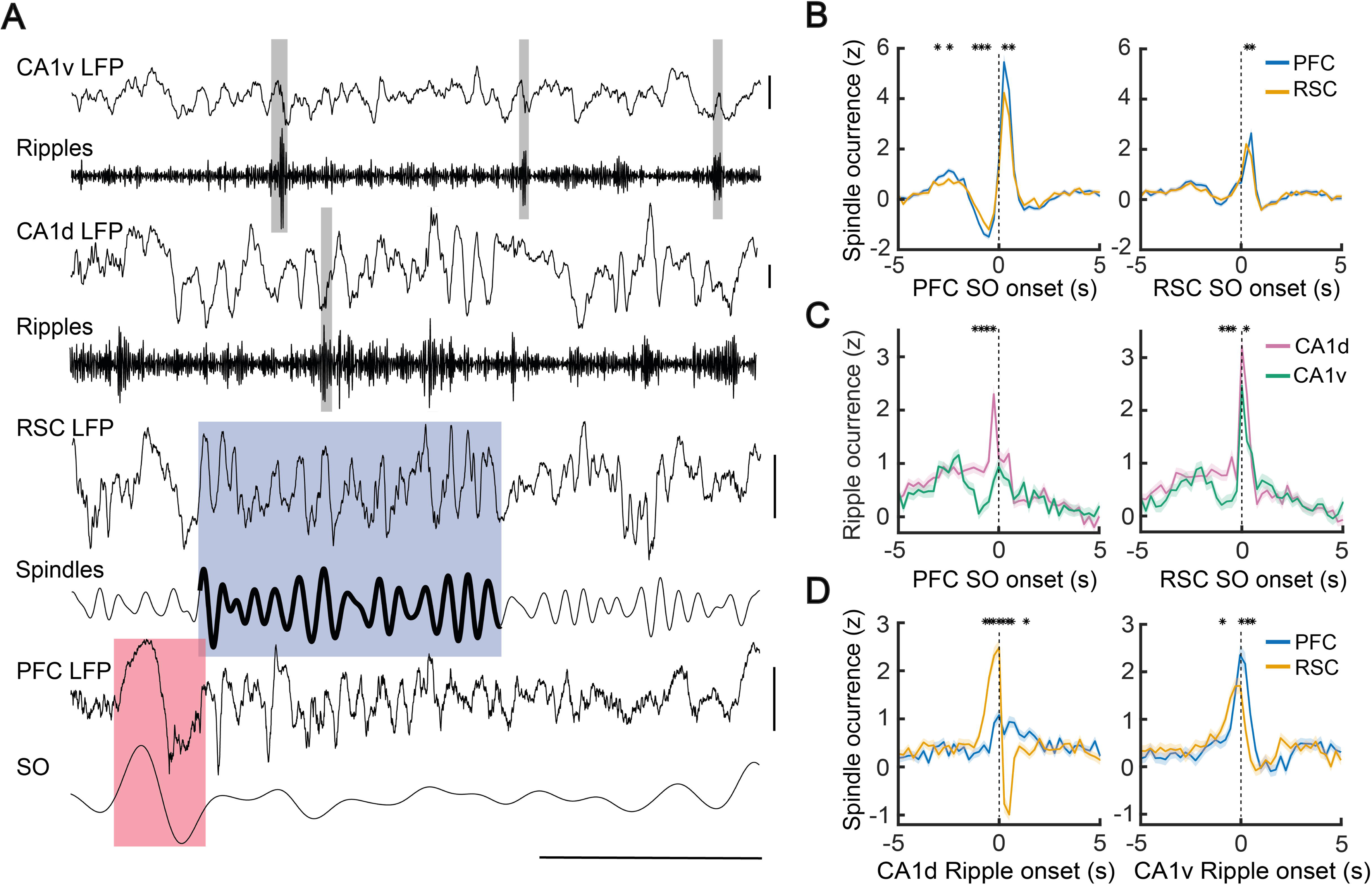
Hierarchical temporal organization of sleep oscillations across hippocampus and neocortex. **A,** Example traces of simultaneously recorded local field potentials (LFPs) from ventral CA1 (CA1v), dorsal CA1 (CA1d), retrosplenial cortex (RSC), and prefrontal cortex (PFC) during nREM sleep (rat PF24, session 9), illustrating canonical sleep oscillatory events. Highlighted segments denote representative ripples in CA1v and CA1d (gray shading), a cortical spindle in RSC (blue shading), and a slow oscillation (SO) in PFC (red shading). Below each LFP trace, the corresponding band-pass filtered signals are shown (ripples: 100–250 Hz; spindles: 7–15 Hz; SOs: 0.1– 4 Hz). Horizontal scale bar: 1 s; vertical scale bar: 0.5 mV. **B,** Temporal coupling of spindle occurrence to SOs in cortex. Z-scored spindle occurrence in PFC (blue) and RSC (orange) aligned to PFC SO onset and RSC SO onset. Spindle latencies were similar between pairs (343.75 ± 7.2 ms, Wilcoxon signed rank test against zero, z = 19.8, p = 2.5×10^-87^). **C,** Temporal coupling of hippocampal ripple occurrence to cortical SOs. Z-scored ripple occurrence in CA1d (magenta) and CA1v (teal) aligned to PFC SO onset and RSC SO onset. Ripple latencies were similar between pairs (138.6 ± 17.7 ms, Wilcoxon signed rank test against zero, z = 6.8, p = 1.2×10^-11^). **D,** Temporal coupling of cortical spindle occurrence to hippocampal ripple onset. Z-scored spindle occurrence in PFC (blue) and RSC (orange) aligned to CA1d and CA1v ripple onset. Spindle latencies were different between pairs (PFC, 332.9 ± 57.1 ms; RSC, −20.7 ± 53.2 ms, Wilcoxon signed rank test, z = 4.9, p = 8.5×10^-7^). In all panels, shaded areas indicate standard error of the mean, and dashed vertical lines mark the onset (t = 0) of the reference event. Asterisks indicate the time bins in which the two traces differ significantly (Wilcoxon signed-rank test across sessions and animals at each bin). P values were corrected for multiple comparisons across all bins using the Benjamini–Hochberg FDR procedure.

### Regional neuronal recruitment by sleep oscillations

Sleep oscillations effectively regulated neuronal firing throughout the hippocampo-cortical circuit (**Table S3**). For example, dorsal ripples simultaneously modulated neuronal discharge locally in the hippocampus (**Fig. 2a**) and distally in the cortex (**Fig. 3a**). Alignment to PFC SO down-state onset revealed distinct regional firing patterns. CA1d exhibited robust triphasic activity, with a pronounced activation preceding the SO (1.99 ± 0.1 z), swift strong suppression during the cortical down-state (−1.91 ± 0.1 z), and a significant posterior rebound (0.63 ± 0.01 z, **Fig. 2b**). Conversely, CA1v showed a weaker response with biphasic modulation, characterized by moderate pre-SO excitation (0.85 ± 0.1 z) and subsequent modest suppression (−0.83 ± 0.1 z), without clear rebound (**Fig. 2b**). All response phases were significantly stronger in dorsal hippocampus (Wilcoxon signed-rank test, FDR-corrected, p < 10^-4^; **Fig. 2b**). Changing the reference to RSC SO maintained a similar, but attenuated triphasic structure in CA1d (peak, 1.72 ± 0.1 z; trough, −0.85 ± 0.1 z; rebound, 0.71 ± 0.1; **Fig. 2b**) and a reduced, biphasic pattern in CA1v (peak, 1.19 ± 0.1 z; trough, −0.68 ± 0.1 z; **Fig. 2b**). Regardless of the SO trigger, dorsal spiking was consistently stronger than ventral neuronal activation (Wilcoxon rank-sum test, z = 4.1, p = 4.82×10^-5^), confirming the larger influence of SOs into the dorsal hippocampus. Cortical regions demonstrated consistent triphasic firing patterns in response to SOs (**Fig. 3b**). PFC SO evoked large spiking triphasic responses in both parietal and frontal neurons, with initial excitation preceding the down-state (PFC, 2.21 ± 0.1 z; RSC, 1.57 ± 0.1 z, Wilcoxon signed-rank test, FDR-corrected, p < 10^-3^), pronounced inhibition at the SO onset (PFC, −4.44 ± 0.3 z; RSC, −2.53 ± 0.2 z; Wilcoxon signed-rank test, FDR-corrected, p < 10^-4^), and clear rebound excitation (PFC, 0.37 ± 0.1 z; RSC, 0.43 ± 0.1 z; Wilcoxon signed-rank test, FDR-corrected, p > 0.05, **Fig. 3b**). Similarly, RSC SO discharged parietal and frontal neurons, with the initial large peak (PFC, 0.70 ± 0.1 z; RSC, 1.36 ± 0.1 z, Wilcoxon signed-rank test, FDR-corrected, p < 10^-4^), followed by marked inhibition at the down-state onset (PFC, −2.35 ± 0.2 z; RSC, −2.82 ± 0.2 z; Wilcoxon signed-rank test, FDR-corrected, p > 0.05), and later rebound (PFC, 0.35 ± 0.1 z; RSC, 0.79 ± 0.1 z; Wilcoxon signed-rank test, FDR-corrected, p < 10^-4^; **Fig. 3b**). Spiking activity triggered by PFC SOs was consistently stronger than that observed following RSC SOs (Wilcoxon rank-sum test, z = 4.5, p = 6.7 × 10^-1^□), confirming that frontal SOs are more effective drivers of cortical neuronal discharge than parietal waves. Neuronal spiking across hippocampo-cortical circuits was strongly modulated by the phase of SOs, with maximal phase-locking occurring around the decaying phase of the up-state (**Fig. S3**). Together, these observations suggest that frontal SOs broadcast globally and preferentially recruit the dorsal hippocampus, whereas parietal SOs remain local but are nonetheless sufficient to entrain both hippocampal poles, again with a dorsal bias.

**Figure 2.**
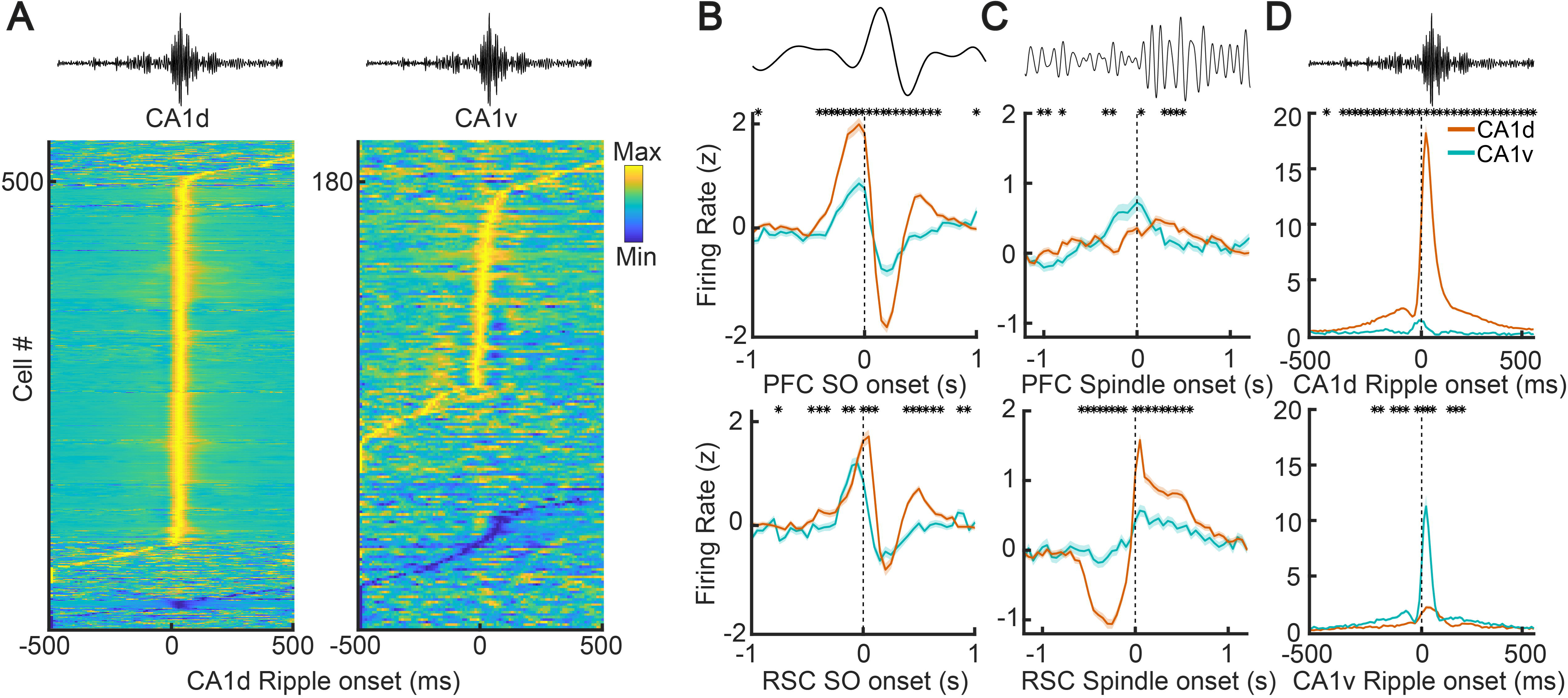
Neuronal recruitment in the hippocampus by sleep oscillations. **A,** Heatmaps show normalized firing rates of individual neurons in dorsal (CA1d, left) and ventral (CA1v, right) hippocampus, aligned to the onset (t = 0 ms) of CA1d ripples. Each row represents the average activity of a single neuron, sorted by response latency, with color indicating firing rate (z-score). **B,** Z-scored mean firing rates of CA1d (orange) and CA1v (teal) neurons aligned to the onset of PFC (top) and RSC (bottom) slow oscillations (SOs). **C,** Normalized firing rates aligned to the onset of PFC (top) and RSC (bottom) spindles. **D,** Normalized firing rates aligned to the onset of CA1d ripples (top) and CA1v ripples (bottom). In all panels, shaded areas represent the standard error of the mean. Dashed vertical lines indicate event onset (t = 0). Asterisks indicate the time bins in which the two traces differ significantly (Wilcoxon signed-rank test across sessions and animals at each bin). P values were corrected for multiple comparisons across all bins using the Benjamini– Hochberg FDR procedure.

**Figure 3.**
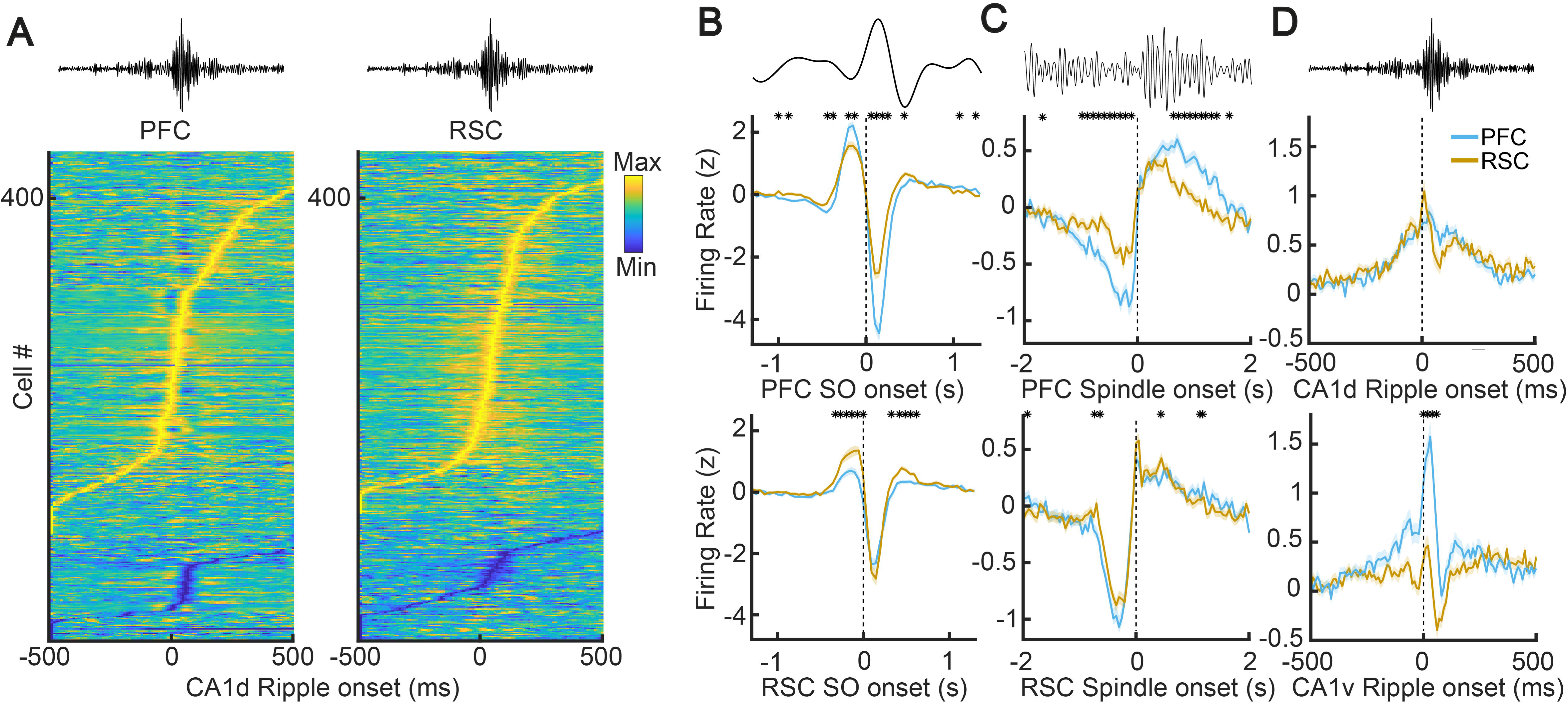
Neuronal recruitment in the cortex by sleep oscillations. **A,** Heatmaps showing normalized firing rates of individual neurons in PFC (left) and RSC (right), aligned to the onset of CA1d ripples (t = 0 ms). Each row represents a single neuron, sorted by response latency, with color indicating firing rate (z-score). **B,** Z-scored mean firing rates of PFC (blue) and RSC (orange) neurons aligned to the onset of PFC SOs (top) and RSC SOs (bottom). **C,** Normalized firing rates aligned to PFC (top) and RSC (bottom) spindle onsets. **D,** Normalized firing rates aligned to the onset of CA1d ripples (top) and CA1v ripples (bottom). Shaded areas represent the standard error of the mean. Dashed vertical lines indicate event onset. Asterisks indicate the time bins in which the two traces differ significantly (Wilcoxon signed-rank test across sessions and animals at each bin). P values were corrected for multiple comparisons across all bins using the Benjamini–Hochberg FDR procedure.

Cortical spindles exerted modest control of neuronal discharge. Frontal spindles elicited small monophasic excitation in both dorsal and ventral hippocampus (CA1d, 0.33 ± 0.1 z; CA1v, 0.28 ± 0.1 z; Wilcoxon signed-rank test, FDR-corrected, p > 0.05, **Fig. 2c**). CA1d displayed contrasting patterns, with pronounced biphasic modulation by parietal spindles (**Fig. 2c**), initially exhibiting significant pre-spindle suppression (−1.08 ± 0.1 z), followed by rebound excitation (1.56 ± 0.1 z). Conversely, CA1v showed exclusively weak monophasic excitation in response to RSC spindles (0.52 ± 0.1 z, Wilcoxon signed-rank test, FDR-corrected, p < 10^-4^, **Fig. 2c**). Thus, cortical spindles weakly influenced the ventral hippocampus. Cortically, spindles triggered biphasic responses in cortical neurons. Indeed, frontal spindles elicited biphasic responses in both PFC and RSC with pre-spindle inhibition (PFC, −0.92 ± 0.1 z; RSC, −0.52 ± 0.1 z; Wilcoxon signed-rank test, FDR-corrected, p < 10^-3^), resulting from the SO down-state preceding spindling, followed by modest activation (PFC, 0.55 ± 0.1 z; RSC, 0.39 ± 0.1 z; Wilcoxon signed-rank test, FDR-corrected, p < 10^-3^; **Fig. 3c**). Similarly, RSC spindles evoked biphasic modulation of both local and distal neuronal discharge with prominent early suppression (PFC, −1.1 ± 0.1 z; RSC, −0.8 ± 0.1 z; Wilcoxon signed-rank test, FDR-corrected, p > 0.05) and modest subsequent excitation (PFC, 0.39 ± 0.1 z; RSC, 0.58 ± 0.1 z; Wilcoxon signed-rank test, FDR-corrected, p > 0.05, **Fig. 3c**). The magnitude and temporal pattern of this response were remarkably similar across regions. Furthermore, cortical spindles organized neuronal firing into distinct oscillation-phases, albeit with weaker modulation than SOs and with effects that varied across regions. (**Fig. S3**). Collectively, spindle oscillations weakly modulate local cortical spiking dynamics (26) and selectively influence hippocampal activity through a preferential dorsal-parietal axis.

Hippocampal ripples differentially controlled neuronal reactivation across the hippocampo-cortical network. Neuronal discharge in CA1d most prominently increased during local dorsal ripples, demonstrating a robust and sharply timed peak (18.14 ± 0.8 z), eliciting a largely minor response in CA1v (1.29 ± 0.1 z, Wilcoxon signed-rank test, FDR-corrected, p < 10^-4^; **Fig. 2d**), probably reflecting residual ripple propagation between hippocampal poles (16). Ventral ripples evoked a similarly selective, yet less prominent, neuronal activation locally in CA1v (11.27 ± 1.0 z), accompanied by a strongly reduced response in CA1d (2.2 ± 0.2 z, Wilcoxon signed-rank test, FDR-corrected, p < 10^-4^; **Fig. 2d**). In cortical regions, ripples generated biphasic responses, albeit with different connection-specific profiles. Dorsal ripples caused significant spiking responses of comparable magnitude in both cortical regions (RSC, 1.04 ± 0.1 z; PFC, 0.84 ± 0.1 z; Wilcoxon signed-rank test, FDR-corrected, p > 0.05; **Fig. 3d**). A different pattern was generated by ventral ripples, which evoked substantial modulation of PFC spiking (1.57 ± 0.2 z), while RSC showed weaker firing modulation (0.46 ± 0.1 z, Wilcoxon signed-rank test, FDR-corrected, p < 10^-4^; **Fig. 3d**). Interestingly, the ventral hippocampus was most effective in engaging cortical neurons, particularly in the frontal cortex (one-way ANOVA F(1,1570) = 22.4, p = 2.3×10^-10^, post-hoc Tukey comparison, p = 2.8×10^-10^). These results reveal a topographic organization during nREM sleep, in which ripple-associated cortical reactivation flows through anatomically constrained pathways, and the dorsal hippocampus emerges as a preferential hub, robustly modulated by the three cardinal sleep oscillations.

### Ripple synchrony gates hippocampo-cortical reactivation

Despite their distinct connectivity profiles and differential dynamics, the dorsal and ventral hippocampal poles exhibited substantial coordination during ripple events (**Fig. 4a**). Hippocampal ripples showed low intrinsic rates (0.53 ± 0.1 Hz) and brief mean durations (57.27 ± 0.1 ms). Under such conditions, if ripples were independently generated in the dorsal and ventral poles, purely random overlap would be expected in a minor fraction of events (about 3%). In contrast, we observed synchronization in a larger proportion of ripples (>12%) between the dorsal and ventral hippocampal poles, suggesting active septo-temporal coordination rather than chance temporal alignment (Wilcoxon signed-rank test against 3%, z = 9.1, p = 7.4×10^-20^). To further investigate intra-hippocampal synchrony and its functional consequences on cortical circuits, we classified sharp wave ripples according to their temporal overlap as independent ripples (CA1d, 84 ± 0.3 % of events; CA1v, 87.7 ± 0.3 % of events) and coordinated CA1d–CA1v ripples (CA1d, 15 ± 0.7% of events; CA1v, 12.2 ± 0.7% of events). As expected by their relative abundance, independent ripples were more frequent than coordinated ripples (Wilcoxon signed-rank test, z = −13.5, p = 1.5×10^-41^, **Fig. S4**). The duration of coordinated ripples was slightly longer than independent events in both hippocampal poles (Wilcoxon signed-rank test, z = 6.1, p = 10^-9^, **Fig. S4**), pointing to intra-hippocampal synchrony as a candidate mechanism regulating the intrinsic structure of ripple episodes. To assess whether ripples were synchronized by propagation across the septo-temporal axis (16), we computed the dorso-ventral ripple crosscorrelogram (**Fig. 4b**). Indeed, when using CA1d as temporal reference, we found that the distribution of ventral onsets was sharply centered at short positive delays (5.90 ± 0.4 ms). Also, the distribution of mean delays indicated that in most cases (77.6%, 83 out of 107 sessions), coordinated ripples spread from dorsal to ventral pole (**Fig. 4b**), also consistent with the larger amplitude of coordinated ripples specifically in the ventral hippocampus (**Fig. S4**). Because cortical SOs can effectively trigger hippocampal ripple events (27), we next analyzed SOs immediately preceding ripples, distinguishing between independent and coordinated ripple episodes. Peak amplitude, onset latency, and total duration of the cortical SOs preceding ripples were comparable between independent and coordinated events (**Fig. S5**). Interestingly, the rising slope of SOs preceding coordinated ripples was modestly yet significantly shallower compared to independent ripples (Wilcoxon rank-sum test, z = −1.6, p < 0.03; **Fig. S5**). Remarkably, frontal firing rates during both SO down and up states preceding coordinated ripples were higher compared to those preceding independent ripple events (Wilcoxon rank-sum test, z = 41.4, p < 10^-6^, **Fig. S5**). These findings suggest that a gradual transition into the SO down-state, combined with increased cortical discharge during SOs, likely enhances the probability of dorsal hippocampal burst propagation ventrally, thus generating a coordinated ripple.

**Figure 4.**
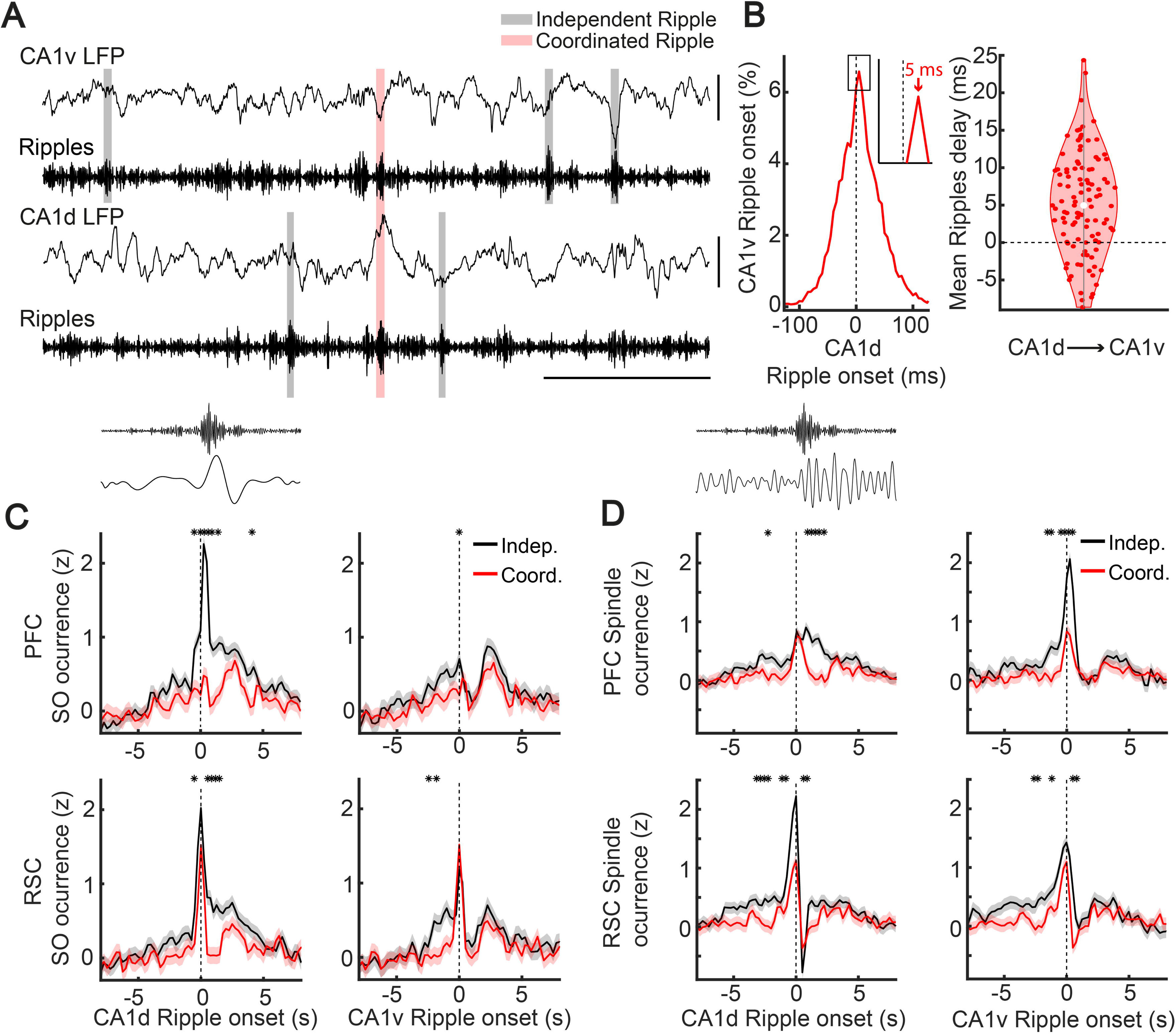
Effect of coordinated versus independent ripples on hippocampo-cortical connectivity. **A,** Example simultaneous LFP recordings from CA1v and CA1d (rat PF40, session 5), showing example independent (gray shading) and coordinated (pink shading) ripple events, with corresponding filtered ripple band signals below. **B,** Left: Distribution of CA1v ripple onset times relative to CA1d ripple onset during coordinated events, indicating the mean delay (6 ms, red arrow). Right: Violin plot of the mean ripple delay for CA1d→CA1v propagation across events (Wilcoxon signed-rank test against 0, z = 6.5, p=7.3×10^-11^). **C,** Z-scored occurrence of slow oscillations (SOs) in PFC (top) and RSC (bottom), time-locked to the onset of independent (black) and coordinated (red) CA1d (left) and CA1v (right) ripples. **D,** Z-scored occurrence of spindle events in PFC (top) and RSC (bottom), time-locked to ripple onset as in (C). In panels C and D, solid lines show the mean, shaded areas represent standard error of the mean, and asterisks indicate time points with significant differences. Dashed vertical lines mark ripple onset (t = 0). Asterisks indicate the time bins in which the two traces differ significantly (Wilcoxon signed-rank test across sessions and animals at each bin). P values were corrected for multiple comparisons across all bins using the Benjamini– Hochberg FDR procedure.

To determine the impact of ripple synchrony on hippocampo-cortical dynamics, we first studied ripple coordination with cortical SOs, yet explicitly segregating ripples into independent and coordinated events. Unexpectedly, this separation uncovered clear interferent cortical dynamics. In fact, coordinated ripple events consistently led to significantly depressed cortical SO associations compared to independent ripples (**Fig. 4c**). Specifically, the cortical SO response typically triggered by CA1d was the most affected when ripples coordinated (Wilcoxon signed-rank test, FDR-corrected, p < 10^-4^, **Fig. 4c**), indicating an attenuated hippocampo-cortical communication. Conversely, CA1v ripples, which generally triggered weaker cortical responses, showed no further reduction in SO activity under dorsal-ventral synchrony (Wilcoxon signed-rank test; FDR-corrected, p < 10^-4^, **Fig. 4c**). Moreover, synchronised ripple events consistently led to significantly weaker spindle responses compared to independent ripples (Wilcoxon signed-rank test; FDR-corrected, p < 10^-4^, **Fig. 4d**). Once again, the effect was more pronounced for the synaptically connected hippocampo-cortical regions. That is, suppression was deeper in CA1d-RSC and CA1v-PFC pathways than in the CA1v-RSC and CA1d-PFC pairs (Wilcoxon signed-rank test; z = 4.2, p = 2.9×10^-5^, **Fig. 4d**). Because ripple propagation along the hippocampal septo-temporal axis was directionally biased, we reasoned that the temporal correlations between sleep oscillations would possibly differ according to such spreading direction within the hippocampus. Nevertheless, regardless of the hippocampal leading pole, spindle-evoked responses in the cortex were strikingly similar (Wilcoxon signed-rank test, FDR-corrected, p > 0.05, **Fig. S6**). Likewise, SO responses in both cortical regions remained nearly identical regardless of ripple propagation direction (Wilcoxon signed-rank test, FDR-corrected, p > 0.05, **Fig. S6**). In addition, no significant differences were apparent in the latencies of evoked responses between independent (191.5 ± 45.6 ms) and coordinated ripples (152.4 ± 55.7 ms, Wilcoxon signed-rank test; z = 0.5, p = 0.6, **Fig. 4d**), suggesting that coordinated and independent ripples engage chiefly the same hippocampo-cortical pathways, yet with different synaptic strength. Thus, cortical spiking activity during the SO, rather than its precise temporal pattern, biases the hippocampus network toward either focal reactivation (i.e.; independent ripples) or septo-temporal synchronization (i.e.; coordinated ripples), connecting cortical dynamics to the emergence of ripple synchrony.

### Brain state modulates ripple-driven reactivation

Remarkably, ripples coordination prominently altered hippocampal spiking patterns. In CA1d, independent ripples elicited the highest local firing peak (15.8 ± 0.9), whereas coordinated ripples produced a significantly smaller local spiking response (5.8 ± 0.4 z; Wilcoxon signed-rank test, FDR-corrected, p < 10^-3^), but simultaneously induced a robust activation in CA1v (1.88 ± 0.2 z), which was absent during independent CA1d events (0.39 ± 0.1 z; Wilcoxon signed-rank test, FDR-corrected, p < 10^-3^, **Fig. 5a**). A mirror pattern emerged for CA1v ripples, where independent events generated the maximal local response in CA1v (10.1 ± 1.1 z), while coordinated ripples resulted in a substantially reduced local response (3.6 ± 0.4 z; Wilcoxon signed-rank test, FDR-corrected, p < 10^-3^), yet recruited CA1d (3.94 ± 0.3 z), which was otherwise modestly activated (0.93 ± 0.1 z; Wilcoxon signed-rank test, FDR-corrected, p < 10^-3^, **Fig. 5a**). Robust activation of the distal pole was observed primarily during coordinated ripples, underscoring the role of active septo-temporal drive in ripple propagation. Notably, neural activation at the distal pole induced by coordinated ripples was consistently smaller than that evoked by independent local events (Wilcoxon signed-rank test, z = 7.8, p = 6.5×10^-15^), suggesting partial recruitment or a reduction in overall excitatory drive during coordinated ripples. This was further supported by the temporal profile of local spiking responses, since the peak response latency remained unchanged between ripple type (independent, 36.19 ± 0.9 ms; coordinated, 38.31 ± 1.0 ms; Wilcoxon signed-rank test, z= −1.6, p = 0.12), while decay time constants were significantly slower for coordinated ripples (independent, −13.7 ± 0.7 s^-1^; coordinated, −6.76 ± 0.5 s^-1^; Wilcoxon signed-rank test; z = −6.5, p = 5.8 × 10^-11^), consistent with their longer duration (**Fig. S4**). These results suggest a diminished excitatory barrage during coordinated events, which likely accounts for the observed depression of local spiking responses. Further analysis of excitation–inhibition dynamics revealed that, despite the overall reduction in activation, the balance between excitatory and inhibitory activity remained stable. Specifically, we classified putative interneurons and pyramidal cells based on action potential waveform features (**Fig. S7**) and calculated the excitatory-to-inhibitory spiking ratio. Excitation increased relative to inhibition during ripple episodes in both dorsal and ventral hippocampus, and this ratio was consistently maintained for both independent and coordinated ripples (**Fig. S8**), indicating that excitation–inhibition balance is preserved during ripple synchrony even when overall excitatory drive is reduced.

**Figure 5.**
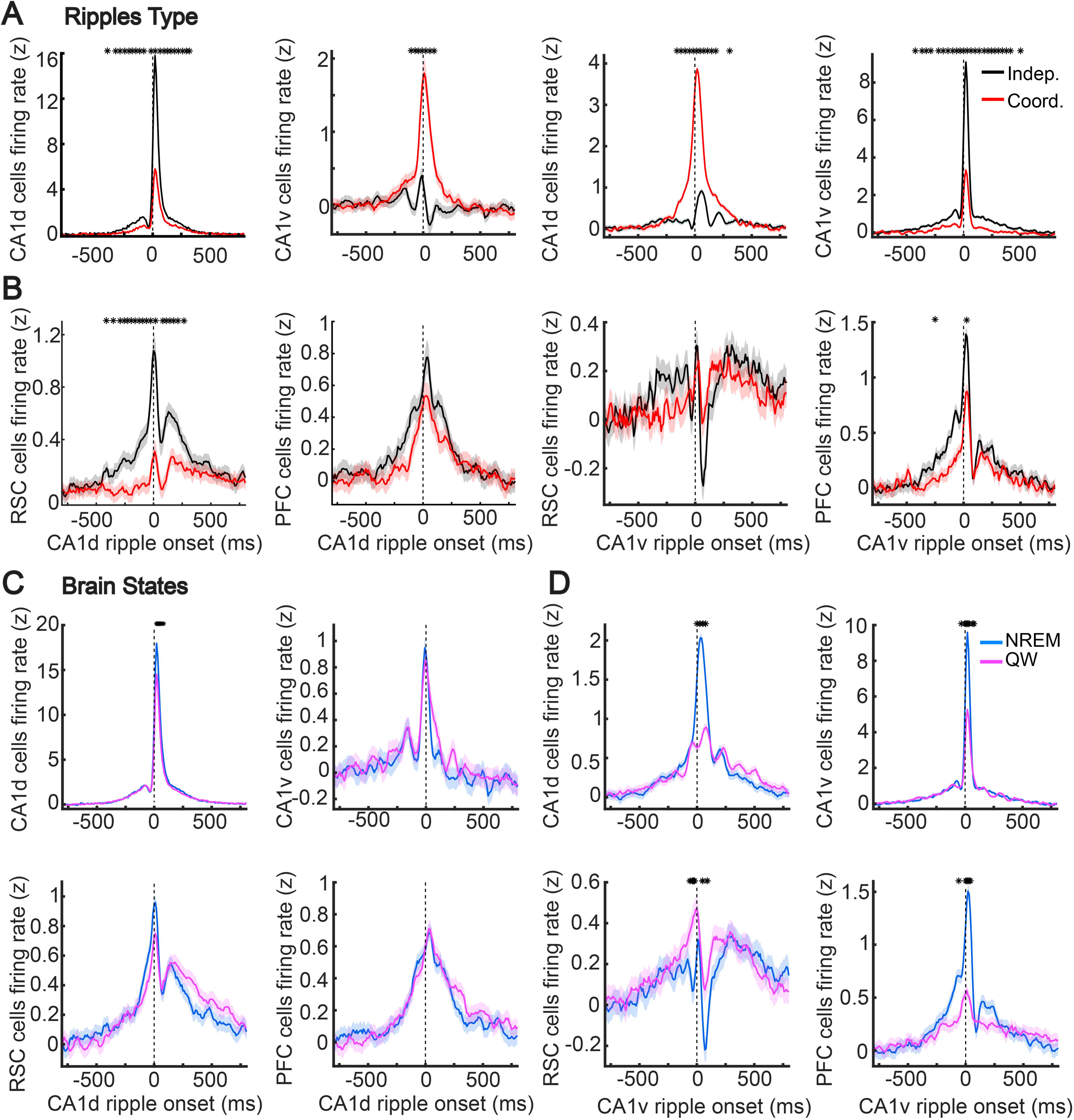
Ripple type and brain state modulation of ripple-triggered neuronal discharge across hippocampal and cortical regions. **A-B,** Mean z-scored firing rates of CA1d and CA1v (A), as well as RSC and PFC (B) neurons, aligned to the onset of independent (black) and coordinated (red) ripple events. **C-D,** Mean firing rates of hippocampal (CA1d and CA1v) and cortical (RSC and PFC) neurons aligned to the onset of CA1d (C) and CA1v (D) ripples during nREM sleep (blue) and quiet wakefulness (QW, magenta). In all panels, shaded areas represent the standard error of the mean, dashed vertical lines mark ripple onset (t = 0). Asterisks indicate the time bins in which the two traces differ significantly (Wilcoxon signed-rank test across sessions and animals at each bin). P values were corrected for multiple comparisons across all bins using the Benjamini–Hochberg FDR procedure.

Next, we evaluated the effect of ripple synchrony on the neuronal spiking of the cortical network. We aligned RSC spiking activity to independent ripples detected exclusively in CA1d and to coordinated dorsal-leading ripple pairs (**Fig. 5b**). Independent CA1d ripples drove a sizable RSC spike burst (1.14 ± 0.1 z), whereas the same cortical population response was significantly suppressed (0.32 ± 0.1 z) when the dorsal ripple propagated to the ventral pole (Wilcoxon signed-rank test; FDR-corrected, p < 10^-4^, **Fig. 5b**). Ripple coordination was ineffective in modifying the PFC spiking profile associated to CA1d. We computed the complementary comparison for PFC, with independent ripples detected only in CA1v. Similarly, coordinated ripples were unable to modulate the modest RSC discharge correlated with CA1v activation. Conversely, CA1v ripples elicited a prominent response in PFC (1.35 ± 0.2 z), which significantly depressed at the peak (0.86 ± 0.1 z) when the ventral ripple propagated dorsally (Wilcoxon signed-rank test; FDR-corrected, p < 10^-4^, **Fig. 5b**). Together, these patterns show that dorsal–ventral synchrony redistributes, rather than augments, hippocampal excitation, sharply dampening local spiking while spreading a weaker volley to the distant pole, suggesting that coordinated ripples act as an intrinsic gate that tempers hippocampo-cortical output.

Contrasting with thalamic spindles and cortical SOs, which are expressed exclusively during sleep, sharp-wave ripples emerge during both nREM sleep and quiet wakefulness (19). To test whether ripple-driven hippocampo-cortical transfer is brain state-dependent, we compared population firing aligned to ripple events recorded in nREM and quiet wake during the same recording sessions (**Fig. 5c-d**). When referenced to CA1d ripples, local neuronal activation remained strong and sharply timed in both states, but was significantly larger during sleep (sleep, 16.56 ± 0.8 z; wake, 13.30 ± 0.7 z; Wilcoxon signed-rank test; FDR-corrected, p <10^-4^, **Fig. 5c**), suggesting potentiation of the dorsal ripple generator (28). Downstream responses were nonetheless brain state-invariant, with CA1v (sleep, 0.95 ± 0.1 z, wake, 0.85 ± 0.1 z; Wilcoxon signed-rank test; FDR-corrected, p > 0.05), RSC (sleep, 0.96 ± 0.1 z, wake, 0.75 ± 0.1 z; Wilcoxon signed-rank test; FDR-corrected, p > 0.05), and PFC (sleep, 0.68 ± 0.1 z, wake, 0.71 ± 0.1 z; Wilcoxon signed-rank test; FDR-corrected, p > 0.05,) showing comparable activation profiles, consistent with the lack of a significant main effect of state (**Fig. 5c**). However, an unanticipated pattern emerged when correlations were triggered by CA1v ripples. Indeed, the local ventral reactivation burst was markedly reduced during wakefulness (sleep, 10.6 ± 1.1 z, wake, 5.68 ± 0.5 z; Wilcoxon signed-rank test; FDR-corrected, p < 10^-4^), and both dorsal hippocampal (sleep, 2.03 ± 0.2 z, wake, 0.89 ± 0.1 z; Wilcoxon signed-rank test; FDR-corrected, p < 10^-4^) and PFC excitation (sleep, 1.51 ± 0.2 z, wake, 0.57 ± 0.1 z; Wilcoxon signed-rank test; FDR-corrected, p < 10^-4^) were also strongly attenuated (**Fig. 5d**). RSC maintained its biphasic response pattern, yet both phases were reduced during wakefulness, the initial activation (sleep, 0.26 ± 0.1 z, wake, 0.47 ± 0.1 z; Wilcoxon signed-rank test; FDR-corrected, p < 10^-4^), followed by the significant suppression (sleep, −0.22 ± 0.1 z, wake, 0.07 ± 0.07 z; Wilcoxon signed-rank test; FDR-corrected, p < 10^-4^, **Fig. 5d**). Thus, ripple-mediated hippocampo-cortical communication is asymmetric across brain states, with dorsal ripples transmitting effectively during sleep and wake, whereas ventral ripples reach cortical targets chiefly during nREM.

Finally, we conducted a comprehensive statistical analysis to evaluate how the multiple factors examined jointly influence neuronal firing during ripple events, thereby illustrating the complex interactions that we previously described individually. Hence, we analyzed neuronal firing rates during hippocampal ripple events using linear mixed-effects models, incorporating ripple type, brain state, ripple initiation pole, and recording region as fixed factors. Animal identity and recording session were included as random effects to account for repeated measures and nested data structures (**Table S4**). Linear mixed-effects modeling revealed that, despite a large number of observations yielding high statistical significance for most main effects and interactions, the experimental factors studied together accounted for only a modest proportion of the variance in neuronal firing rates (fixed-effects marginal R² = 0.052, conditional R² = 0.058, **Table S5**). Variance decomposition indicated that animal and session random effects accounted for less than 1% of the total variance, with the majority arising from within-session (trial-to-trial) variability (**Table S5**). After obtaining the parameter estimates, we performed an ANOVA marginal test to determine the fixed effects significance (**Table S6**). Interestingly, the most significant effect was yielded by the ripple type*ripple initiation pole interaction (p = 3.82×10^-54^). Moreover, robust main effects of recording region (p = 7.3×10^-26^), ripple hippocampal pole (p = 1.4×10^-5^), and especially ripple type (p = 7.0×10^-43^) were observed, along with significant higher-order interactions (**Table S6**), such as recording region*ripple type*ripple initiation pole (p = 9.89×10^-50^, **Fig. 5a**). Importantly, the effect of brain state alone was not significant (p = 0.16), yet brain state contributed to several significant interactions, such as recording region*brain state*ripple initiation pole (p = 2.64×10^⁻15^, **Fig. 5c**).

Collectively, these findings demonstrate that ripple-driven neuronal recruitment across hippocampo-cortical networks is shaped by intricate interactions involving several factors, underscoring complex mechanisms underpinning memory-related hippocampo-cortical communication. Among those factors, ripple synchronization emerges as a key determinant of the spike timing across hippocampal and cortical circuits.

## DISCUSSION

Memory consolidation fundamentally depends on the orchestrated interaction of canonical sleep oscillations: cortical slow oscillations, thalamo-cortical spindles, and hippocampal sharp-wave ripples. Yet, the organizational logic and the functional rules guiding this cross-structure dialogue remain only partially understood. Our results provide a hierarchical and mechanistically nuanced view of how these oscillatory events coordinate the flow of information within the hippocampo-cortical network during memory consolidation. Specifically, we reveal that SOs originating in frontal cortex serve as a global temporal scaffold, resetting thalamic circuits and launching spindle volleys that propagate from anterior to posterior across the cortex, consistent with prior findings in both rodents and humans (2, 29, 30). Spindles, in turn, act as region-specific channels, with parietal spindles triggering robust, biphasic suppression– rebound modulation of dorsal hippocampus, while frontal spindles provide weaker, monophasic excitation that influences both hippocampal poles. This regional selectivity establishes anatomically matched feedback loops, which set the stage for precisely targeted memory reactivation.

Our work extends the active systems consolidation model (4, 5) by showing that hippocampo-cortical interactions are not simply global synchronizations but are routed through distinct anatomical pathways, forming a multilayered network in which specificity and selectivity are paramount. In this architecture, hippocampal ripples act as content carriers whose cortical targets mirror their septo-temporal origin, as dorsal ripples preferentially recruit the parietal cortex, whereas ventral ripples more broadly engage both frontal and parietal regions. This pole-specific recruitment aligns with gene expression and anatomical data suggesting a blend of discrete functional domains and continuous gradients along the hippocampal long axis (6, 31). Such anatomical specificity supports the idea that different hippocampal subregions may convey spatial versus emotional and motivational content, consistent with the well established role of dorsal hippocampus in spatial navigation and the ventral pole involvement in affective processing (9, 32).

A central finding of our study is that synchrony between dorsal and ventral hippocampal poles during ripple episodes does not amplify but rather suppresses cortical reactivation. Indeed, when ripples occur simultaneously in both hippocampal poles, a phenomenon we observed to be more frequent than expected by chance, local excitatory drive at the initiating pole is decreased and redistributed rather than globally enhanced. This is evident as a marked reduction in local neuronal discharge and a collapse of spindle and SO responses in the cortex, despite preserved ripple waveform and unchanged excitatory-inhibitory balance at the source. These results challenge the widespread assumption that increased hippocampal synchrony universally strengthens memory replay and instead reveal that dorso-ventral ripple coordination acts as a dynamic gate (24, 33). Rather than broadly promoting replay, this gating mechanism ensures that only the most relevant memory traces, likely those with the strongest initial activation, are integrated into cortical networks at high gain, while broader, lower-gain coordinated events suppress nonspecific or potentially interfering reactivation.

Our data further highlight that the microstructure of preceding cortical SOs can bias the hippocampal network toward either local, high-gain replay or septo-temporally coordinated, lower-gain events. This finding supports a model in which slow waves and their transitions not only group spindles and ripples within up-states (34, 35) but also dynamically gate the specificity of hippocampal output to cortex. This is also consistent with recent observation that dCA1 and RSC operate as weakly coupled excitable systems capable of bidirectional perturbation, with the RSC potentially serving as a gateway that enables hippocampal ripples to propagate SO down-states to downstream cortical regions via cortico-cortical pathways (36). This alternation between up- and down-states may reflect alternating windows for active consolidation and homeostatic recalibration (37). The PFC, in particular, exhibits pronounced firing suppression during SO down-states, aligning with its proposed role as a site of long-term associative memory storage and plasticity (38).

We also show that ripple-driven communication is strongly modulated by brain state, but primarily through its interactions with other factors. For example, dCA1 ripples are effective in driving cortical targets during both sleep and quiet wakefulness, whereas vCA1 ripples preferentially amplify hippocampo-cortical coupling during nREM sleep. This is consistent with work demonstrating privileged reactivation of hippocampal ensembles during sleep (39, 40) and suggests that nREM may provide optimal network conditions for the integration of emotionally or motivationally significant information, often processed via the ventral hippocampus (41). We thus propose a dual-level control, in which SO phase and brain state act globally, while ripple synchrony provides a local, dynamic gating mechanism for hippocampal-cortical transfer.

Overall, these insights refine the active systems consolidation framework, moving beyond simple models where higher synchrony entails stronger replay. Instead, our results argue that hippocampal synchrony operates as a selective brake, protecting the cortex from indiscriminate activation. This dynamic gating allows the hippocampo-cortical network to flexibly prioritize high-value traces, likely spatial in dorsal pathways, and emotional-motivational in ventral connections, and to suppress potentially spurious activation. By mapping how SOs establish a global temporal scaffold, spindles specify which hippocampal pole engages with cortical targets, and ripple synchrony ultimately determines the effectiveness of hippocampo-cortical connectivity, we provide a functional anatomical blueprint for memory consolidation. This is conceptually consistent with findings from large-scale neural population studies, which show that experimentally induced or natural variability often explains only a modest proportion of neural firing rate variance, with the majority attributable to intrinsic and trial-level fluctuations (42–44). In our data, linear mixed-effects modeling confirmed that fixed experimental factors and their interactions explain approximately 5% of the total variance in firing rates, with the remainder reflecting trial-by-trial variability and minor (< 1%) contributions from animal and session identity. This highlights the importance of effect size and context, not just statistical significance, when interpreting large-scale neural datasets (45–47).

By integrating the hierarchy of oscillatory interactions, our findings help to shed light on previous discrepancies regarding the functional significance of hippocampal synchrony and extend the systems consolidation model to account for dynamic, context-dependent gating. Indeed, the selective amplification of hippocampo-cortical axes during sleep may explain the preferential consolidation of emotional or motivational memories observed in some behavioral studies (5, 48). Our approach underscores the necessity of hierarchical statistical modeling and transparent reporting of explained variance to ensure mechanistic interpretations are both meaningful and generalizable. Looking forward, future studies could employ simultaneous recordings from upstream inputs (e.g., CA3 or entorhinal cortex) and targeted manipulations, such as optogenetic silencing or chemogenetic modulation, to delineate the cellular and circuit mechanisms underlying ripple-driven gating (49). Single-cell imaging approaches might further reveal how specific neuronal ensembles participate in or are excluded from cortical reactivation during coordinated versus independent ripple events, providing insight into the microcircuit logic of this dynamic gate (20).

In sum, our findings support a model of memory consolidation in which cortical slow oscillations orchestrate precise ripple–spindle coupling dynamics, hippocampal synchrony acts as a context-sensitive gate for cortical engagement, and the interplay of anatomical cortical specificity, brain state, and local network configuration ensures selective, efficient, and functionally meaningful integration of memory traces. This multilayered control mechanism provides a flexible substrate for sculpting the precision and content of systems consolidation, fundamentally reshaping our understanding of hippocampo-cortical communication during sleep and wake.

## Materials and Methods

Twenty adult male Sprague-Dawley rats (P40–60) were used in accordance with institutional ethical guidelines. Animals were acclimated to the housing environment, handled, and further habituated to the recording setting prior to surgery. Custom 16- or 32-channel recording drives were assembled and stereotaxically implanted to target CA1d, CA1v, RSC, and PFC, with electrode positions verified post hoc by histology. Following recovery, 10 days of electrophysiological recordings were performed in a Faraday-shielded enclosure, with rats monitored during 60–90-minute sessions. Local field potentials (LFP) and single-unit/multiunit activity were acquired at 20 kHz, synchronized with video for behavioral scoring. Brain states (quiet wake, nREM, REM) were classified by LFP patterns, spectrograms, and behavior. Ripple, spindle, and slow oscillations were detected using bandpass filtering and amplitude thresholding with custom MATLAB scripts. Spike sorting was performed using Kilosort2 and manual curation; putative pyramidal cells and interneurons were identified by waveform and firing rate characteristics. At the end of the study, electrode locations were verified by electrolytic lesions and Nissl staining. Statistical analyses included parametric or non-parametric tests as appropriate, repeated-measures ANOVA, post-hoc Bonferroni corrections, and effect size estimation, with significance set at α = 0.05. Analyses were performed in MATLAB.

## ACKNOWLEDGMENTS

This work was supported by grants fondecyt 1230589, Anillos ACT 210053, Redes 190045.

## Author contributions

PF designed research and wrote the manuscript; NE performed most of the analysis with input from PF. GL performed experiments and analysis. MC, GF, AA and AL-V performed experiments.

The authors declare no competing interest

