## Supplemental information for "Hippocampal Synchrony Dynamically Gates Cortical Connectivity Across Brain States"

**Supplementary Information**

**MATERIALS AND METHODS**

**Animals**. Twenty adult male Sprague-Dawley rats (postnatal days 40–60, weighing 300–350 g) were obtained from the Center for Innovation on Biomedical Experimental Models (CIBEM, <https://cibem.bio.puc.cl/>) at the Pontificia Universidad Católica de Chile. All experimental procedures were approved by the Scientific Ethical Committee for the Care of Animals and the Environment (CEC-CAA, protocol 220512003). Rats were housed in a temperature-controlled room (22 ± 1 °C) under a 12-hour light/dark cycle (lights on at 7:00 a.m.) with food and water provided ad libitum.

**Habituation**. Animals were habituated to the housing environment for three days, then handled daily by the experimenter for three to five additional days. Subsequently, rats were habituated to the experimental environment by placing them on a towel-covered drum in the recording room for 30–60 minutes daily over approximately one week.

**Recording Implant Assembly**. The recording drive was designed using Autodesk Fusion software and 3D-printed on a Fusion360 F410 printer. Sixteen or 32 tungsten standard tapered-tip electrodes (MicroProbes, USA; impedances of 1, 3, and 5 MΩ) were assembled onto a custom-fabricated drive targeted to brain regions of interest based on a rat brain stereotaxic atlas (44). Each electrode was connected to an EIB-18 or EIB-36 PCB card (capacity of 18 or 36 channels, including ground and reference). Ground and reference channels were connected to stainless steel screws placed into the skull during surgery. The entire assembly was covered with copper mesh to minimize electrical noise.

**Stereotaxic Surgery**. Rats were anesthetized with isoflurane (4% induction, 1.5–2% maintenance) and secured in a stereotaxic frame (Stoelting Inc.). Body temperature was maintained at 35–37 °C using a homeothermic blanket, and animals received hourly hydration with glucosaline solution (0.9% NaCl, 2.5% dextrose). Following a scalp incision, four craniotomies (~1 mm diameter each) were performed in the right hemisphere at predetermined stereotaxic coordinates (CA1d: -4.0 AP, 2.0 ML, 2.7 DV; CA1v: -5.7 AP, -5.5 ML, 7.2 DV; RSC: -4.0 AP, 0.7 ML, 2.2 DV; PFC: 2.5 AP, -0.7 ML, 4.2 DV, all in mm). Two additional craniotomies anterior to bregma accommodated the ground and reference electrodes, while four more craniotomies (two parietal contralateral, one parietal ipsilateral, and one posterior to lambda) secured the implant with additional screws. The dura was carefully removed in the craniotomies, cortical surfaces were moistened with mineral oil, and electrodes were carefully inserted. Craniotomies were sealed with silicone elastomer or wax, and the recording drive was fixed using dental acrylic. Postoperative care involved daily subcutaneous injections of enrofloxacin (10 mg/kg) and meloxicam (1 mg/kg) for three consecutive days. Animals recovered for at least seven days before recordings began.

**Electrophysiological Recordings**. Following recovery, electrophysiological recordings were conducted over 10 consecutive days during the light phase in a Faraday-shielded enclosure. Rats were placed on a towel-covered platform for 60–90-minute sessions. The EIB-18 or EIB-36 boaard was connected via a 16- or 32 channel headstage (Intan Technologies) to an amplifier (Intan RHD Recording System, Intan Technologies, USA), with one electrode serving as reference. Video recordings synchronized with the amplifier clock aided in brain state identification. Signals were sampled at 20 kHz using RHX Data Acquisition software (Intan Technologies) and later converted to MATLAB format using the LAN toolbox for further analysis.

**Histology**. After the recording protocol finished, rats were anesthetized with isoflurane (4% induction, 1.5–2% maintenance), and electrolytic lesions (5 µA for 10 s) marked electrode locations. Following 48 hours of recovery, animals were terminally anesthetized with ketamine (300 mg/kg) and xylazine (30 mg/kg, i.p.) and transcardially perfused with 0.9% saline followed by 4% paraformaldehyde. Brains were postfixed overnight, transferred to PBS-azide, and sectioned coronally using a vibratome (World Precision Instruments, USA). Sections were Nissl-stained and examined with a Nikon Eclipse CI-L microscope to verify electrode placements.

**Brain State Identification**. We used a previously described method (45, 46) Briefly, brain states (quiet wakefulness, nREM sleep, and REM sleep) were identified via visual inspection of raw LFP from the CA1d region, spectral analysis (Fourier spectrogram), and behavioral observation from video recordings. States were classified in 10-second epochs: quiet wakefulness exhibited mixed-frequency activity (>20 Hz) with minimal movement; nREM sleep displayed continuous low-frequency activity (0.1–4 Hz) without movement; REM sleep showed pronounced theta activity (5–10 Hz) and muscle atonia. State scoring employed custom MATLAB scripts.

**Sleep Oscillations Detection**. Ripples were identified by band-pass filtering LFP (100–250 Hz, zero-phase shift non-causal filter, 0.5 Hz roll-off), rectifying, and low-pass filtering at 20 Hz (4th order Butterworth). Signals were z-score normalized, and events exceeding a 3.5 SD threshold (CA1d) or 2.5 SD (CA1v) lasting ≥50 ms were identified and visually verified. Onset and offset time-points were determined by the crossing of a 1 SD threshold, with a 50 ms refractory period to prevent duplicates. Spectral analysis using 7-cycle Morlet wavelets (adapted from a previously used method (20)) were used to determine ripple frequency, amplitude, and duration. Spindles were identified using LFP band-pass filtering (7–15 Hz, 4th order Butterworth), root mean square envelope smoothing (100 ms Gaussian), and segments exceeding 1 SD for ≥500 ms (adapted from a previously used method (47)). Slow oscillations were isolated by low-pass filtering LFP (0.1–4 Hz, 2nd and 5th order filters, respectively). Oscillations were identified by zero crossings, with thresholds set for positive peaks (>85th percentile) and negative troughs (<40th percentile). Putative slow oscillations were further filtered to periods of nREM sleep (adapted from previously used methods (24, 48)).

**Spike Sorting**. Extracellular signals were band-pass (600-5000 Hz) filtered and the spike waveforms with either negative or positive peaks exceeding were extracted. Single units were sorted using a custom manual clustering program (Kilosort2, <https://github.com/MouseLand/Kilosort>) and were distinguished based on principal components. Putative pyramidal cells and interneurons were further separated by principal component analysis of the unit waveforms and mean firing rat. A unit cluster was classified as a single-unit activity if the refractory period (time between two consecutive spikes) was at least 1.5 ms. When the recording quality and the spike sorting did not allow unambiguous single-unit isolation, the spike cluster was conservatively classified as multiunit activity (MUA).

**Statistical Analysis**. Group differences with a single, independent factor were examined with a one-way ANOVA when the data met parametric assumptions (normality and homogeneity of variances); if normality was violated the non-parametric Kruskal–Wallis test was substituted. For repeated observations, a repeated-measures ANOVA was applied when residuals were normally distributed. If only two repeated levels were compared, a paired-samples *t*-test was used; when the normality assumption was violated a Wilcoxon signed-rank test (two levels) or a Friedman test (≥ 3 levels) replaced the parametric test. Sphericity was assessed with Mauchly’s test, and the Greenhouse–Geisser correction was applied when sphericity was violated. Post-hoc pairwise comparisons after any omnibus ANOVA were adjusted with the Bonferroni procedure. Linear relations between two continuous, normally distributed variables were assessed with Pearson’s *r*; otherwise Spearman’s ρ was used. The threshold for statistical significance was set at two-tailed *α* = 0.05. Effect sizes were reported for all main tests (Cohen’s *d* for *t*-tests, partial η² for ANOVAs, and *r* or ρ for correlations); values of Cohen’s *d* ≥ 0.3 were interpreted as a significant effect. The decay time constant (*b*) was calculated by fitting the data to a single-term exponential decay model of the form *y=a*exp(b*t)* using least-squares regression. The fitting and statistical analyses were performed using the MATLAB software package (MathWorks).

**Supplementary Figures**

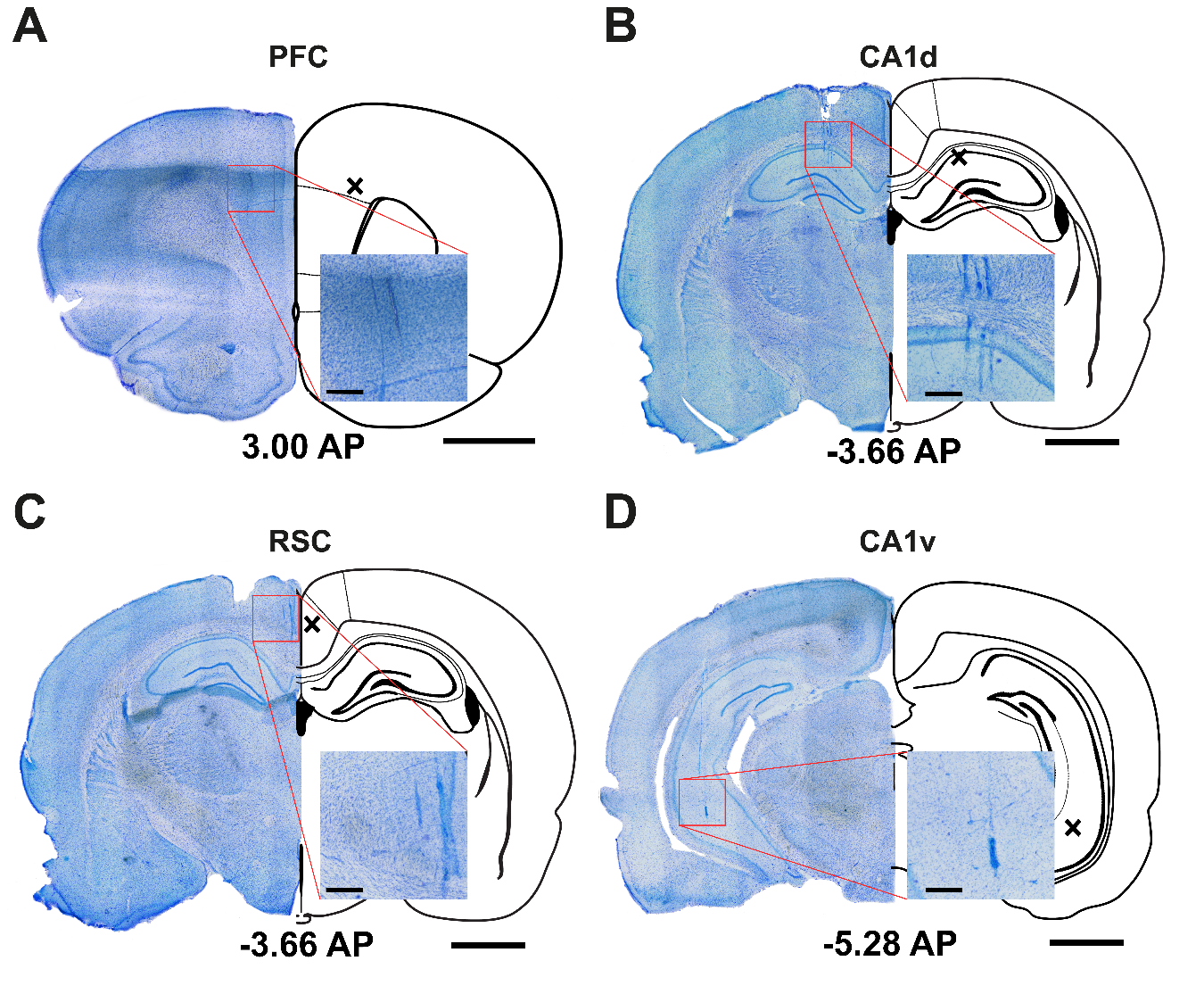

**Figure S1. Histological verification of recording sites.** Representative Nissl-stained coronal brain sections (100 um thick, rat PF07) illustrate electrode tracks and lesion sites at the indicated anterior-posterior (AP) coordinates relative to bregma. **A**, PFC (+3.00 mm AP); **B**, CA1d (−3.66 mm AP); **C**, RSC (−3.66 mm AP); and **D**, CA1v (−5.28 mm AP). Black ‘X’ marks on schematic overlays indicate targeted coordinates for electrode implantation. Insets highlight the electrode tracks or lesions within the corresponding target regions at higher magnification. Scale bars: 1 mm (whole section); 200 µm (inset).

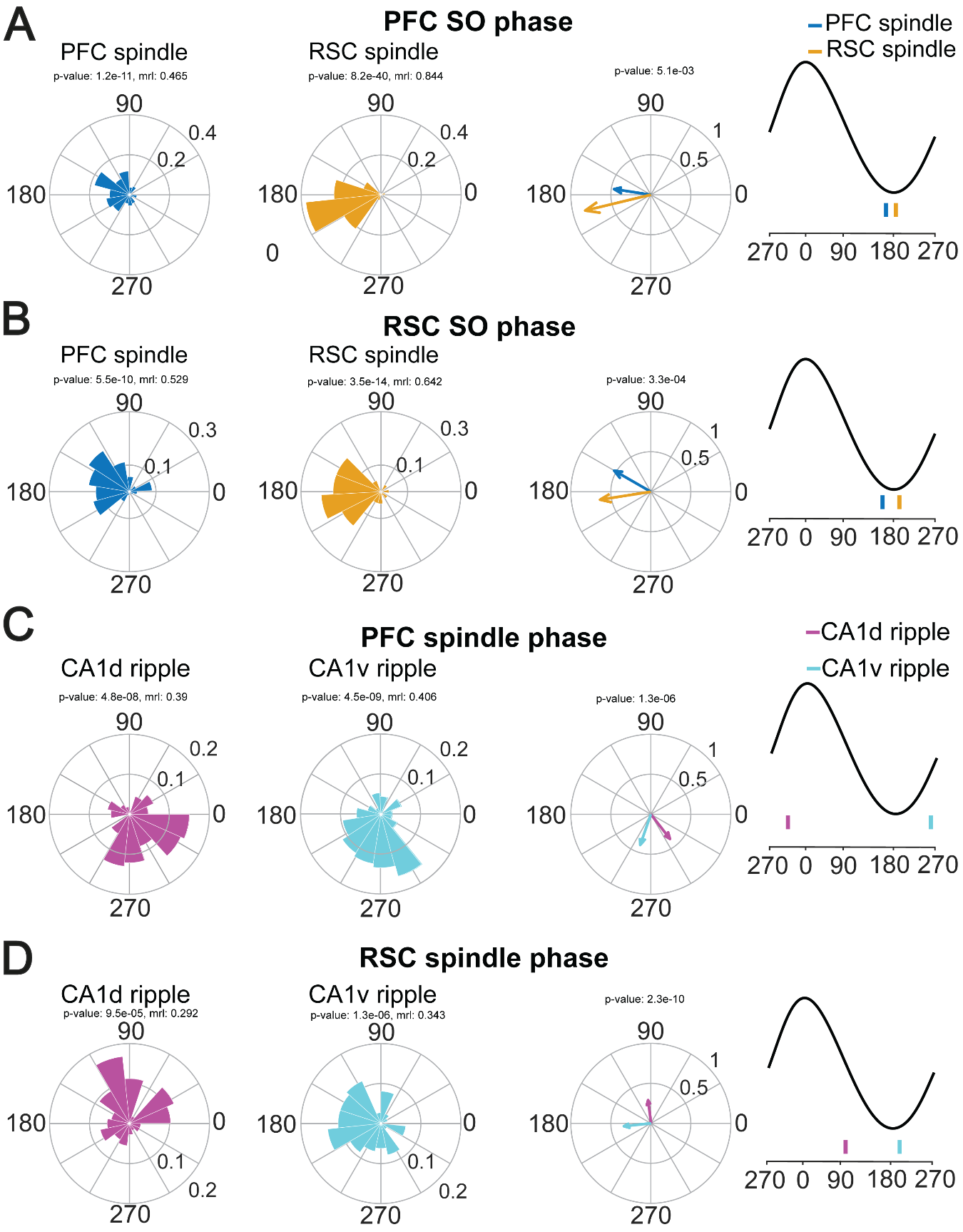

**Figure S2. Cross-regional phase modulation of sleep oscillations and ripple events.** Polar plots illustrate the preferred phase of spindle and ripple event occurrence relative to ongoing SO and spindle cycles in PFC, RSC, CA1d, and CA1v. **A** and **B**, spindle onset probability as a function of SO phase in PFC (blue) and RSC (orange), showing consistent phase-locking to the SO up-state with significant differences in mean phase angle (Rayleigh test, p-values indicated). **C** and **D**, probability of CA1d (magenta) and CA1v (cyan) ripples as a function of spindle phase in PFC and RSC, demonstrating significant, though modest, phase modulation (p-values indicated). Rightmost panels depict the mean phase relationships on a schematic oscillation cycle, with vertical bars indicating the mean preferred phase for each event type and region. MRL: mean resultant vector length.

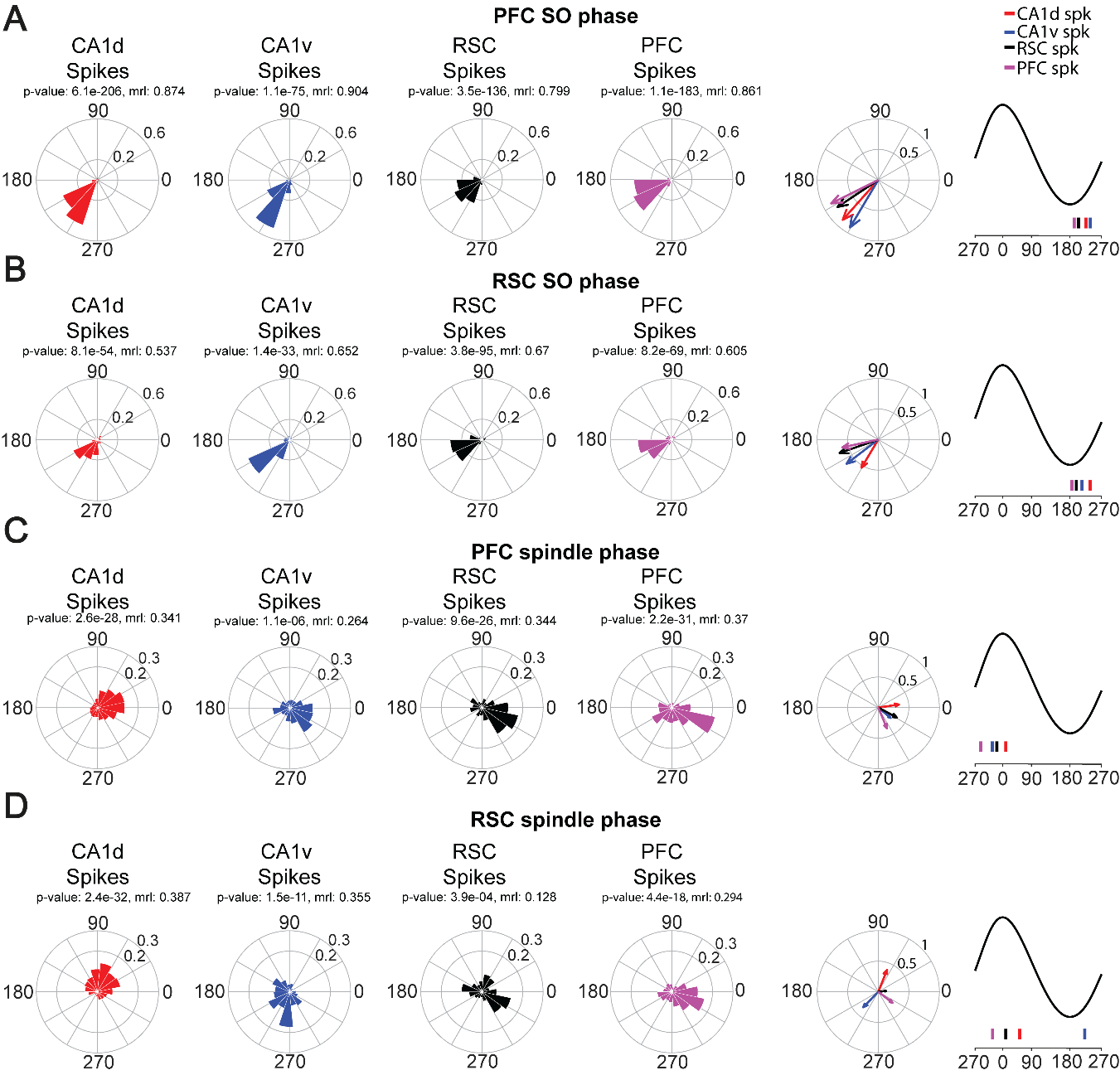

**Figure S3. Preferred phase of neuronal spiking relative to ongoing sleep oscillations across hippocampal and cortical regions during NREM sleep.** Polar plots show the probability distribution of spike timing for CA1d (red), CA1v (blue), RSC (black), and PFC (magenta) neurons as a function of ongoing SO phase in PFC (**A**) or RSC (**B**). Same format as above, but for spike timing relative to the phase of spindle oscillations detected in PFC (**C**) or RSC (**D**). Rightmost panels in each row summarize the mean preferred phases for each population, plotted on the corresponding oscillatory cycle, with colored tick marks denoting the circular mean for each region. For each plot, the Rayleigh p-value and mean resultant vector length (MRL) are indicated.

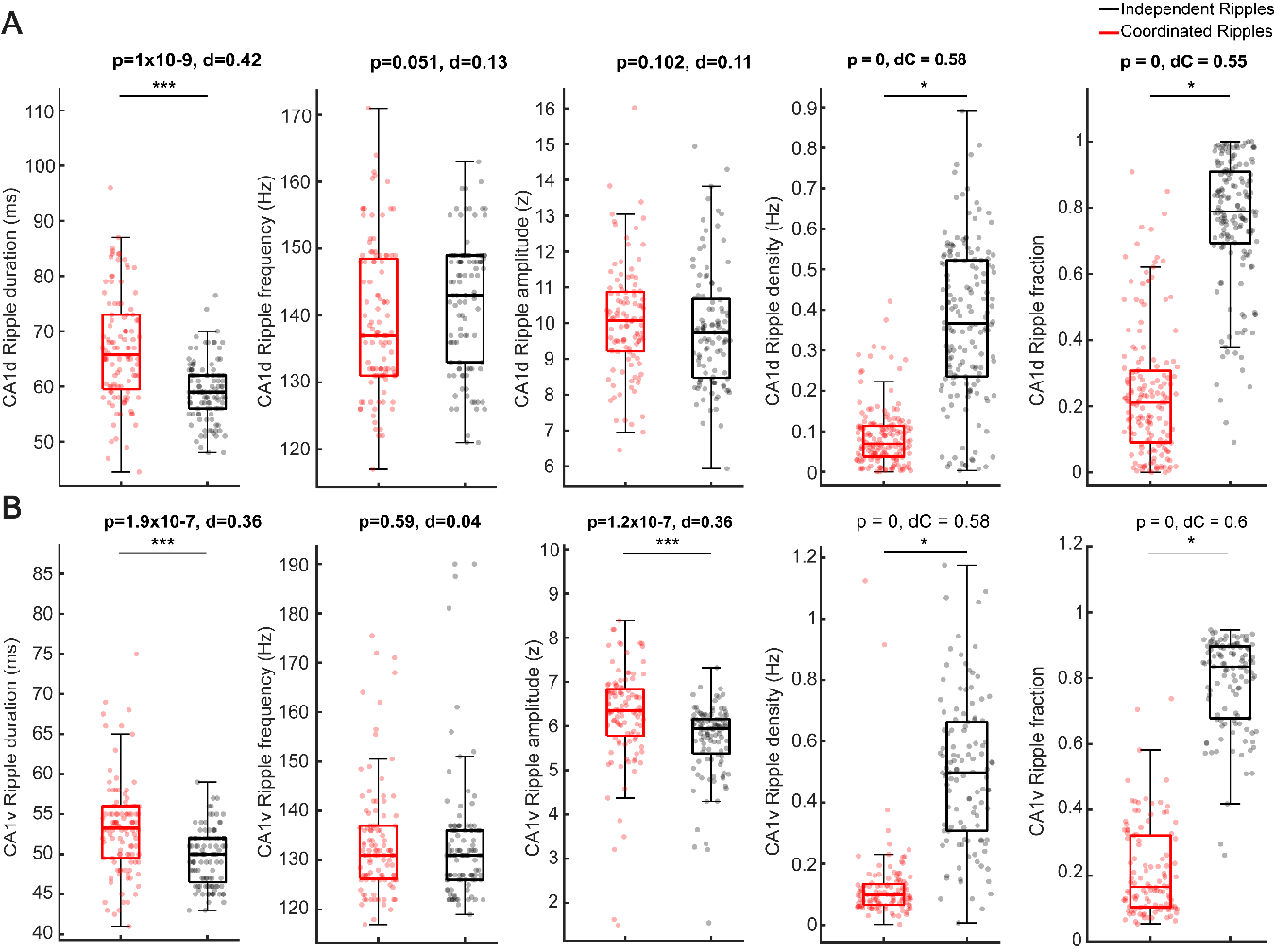

**Figure S4. Intrinsic characteristics of independent and coordinated hippocampal ripples.** Boxplots comparing ripple duration, intra-ripple frequency, amplitude, density, and fraction for coordinated (red) and independent (black) ripples recorded in CA1d (**A**) and CA1v (**B**). In each plot, the box indicates the interquartile range (25th–75th percentile), the horizontal line marks the median, whiskers depict lower (Q1) and upper (Q4) quartiles, and individual ripple events are shown as semi-transparent dots. Statistical comparisons were performed using two-sided Wilcoxon rank-sum tests; p-values and effect sizes (Cohen’s d), d) are indicated above each plot. Asterisks (***) denote highly significant differences (p < 0.001).

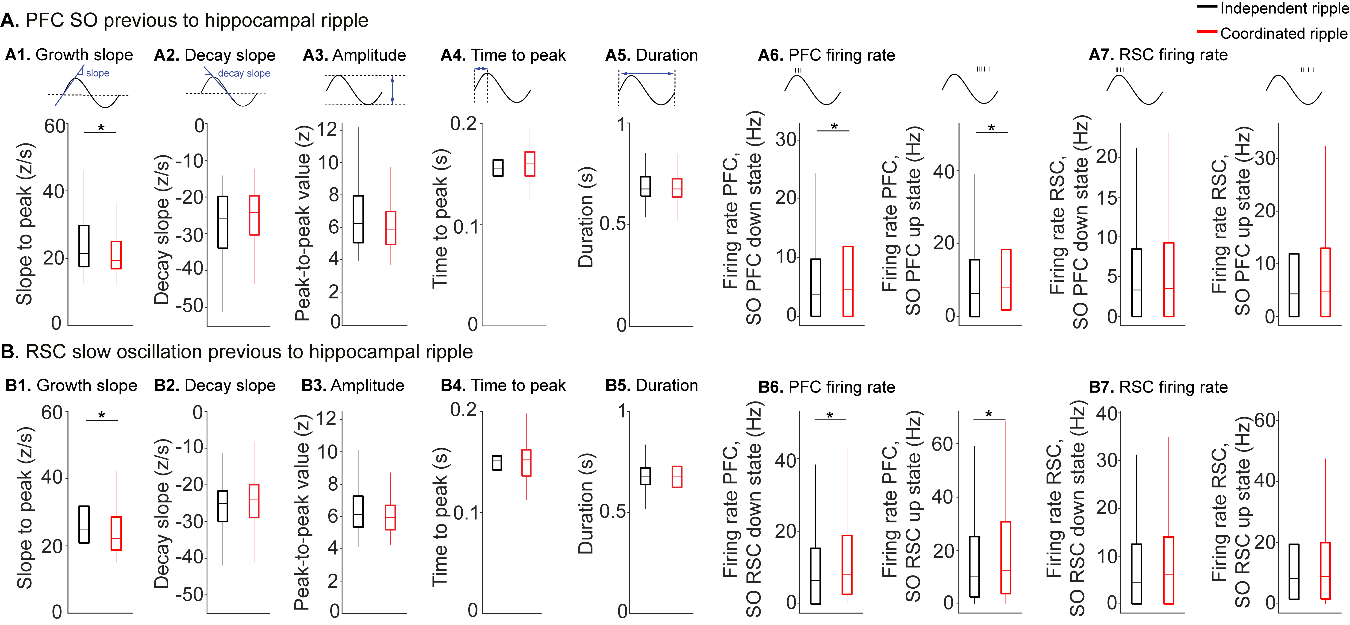

**Figure S5. Comparison of pre-ripple slow oscillation characteristics and firing rates for independent versus coordinated ripples.** Boxplots show features of PFC (**A**) and RSC (**B**) SOs preceding independent (black) and coordinated (red) hippocampal ripples. Panels show SO waveform characteristics: growth slope (A1, B1), decay slope (A2, B2), amplitude (A3, B3), time to peak (A4, B4), and duration (A5, B5). The final panels show neuronal firing rates during the source SOs down- and up-states, separated into local and cross-regional activity. Local firing rates are shown for the PFC during its own SO (A6) and the RSC during its own SO (B7). Cross-regional modulation is shown by the RSC firing rate during the PFC SO (A7) and the PFC firing rate during the RSC SO (B6). In each plot, the box indicates the interquartile range (25th–75th percentile), the horizontal line marks the median, and whiskers depict the rest of the data distribution. Statistical significance was determined by two-sided Wilcoxon rank-sum tests, and effect sizes were calculated using Cohen's d. were observed in the growth slope of both PFC (A1, p = 0.02, d = 0.12) and RSC (B1, p = 4.8x10^-4^, d = 0.20) SOs. firing rates were significantly higher preceding coordinated ripples for the PFC during its own SO down-state (A6 left, p = 7x10^-104^, d = 3.5) and up-state (A6 right, p = 2.1x10^-83^, d = 3.5), and for the PFC firing rate during RSC SO down-state (B6 left, p = 2.4x10^-103^, d = 3.3) and up-state (B6 right, p = 2.4x10^-66^, d = 3.7).

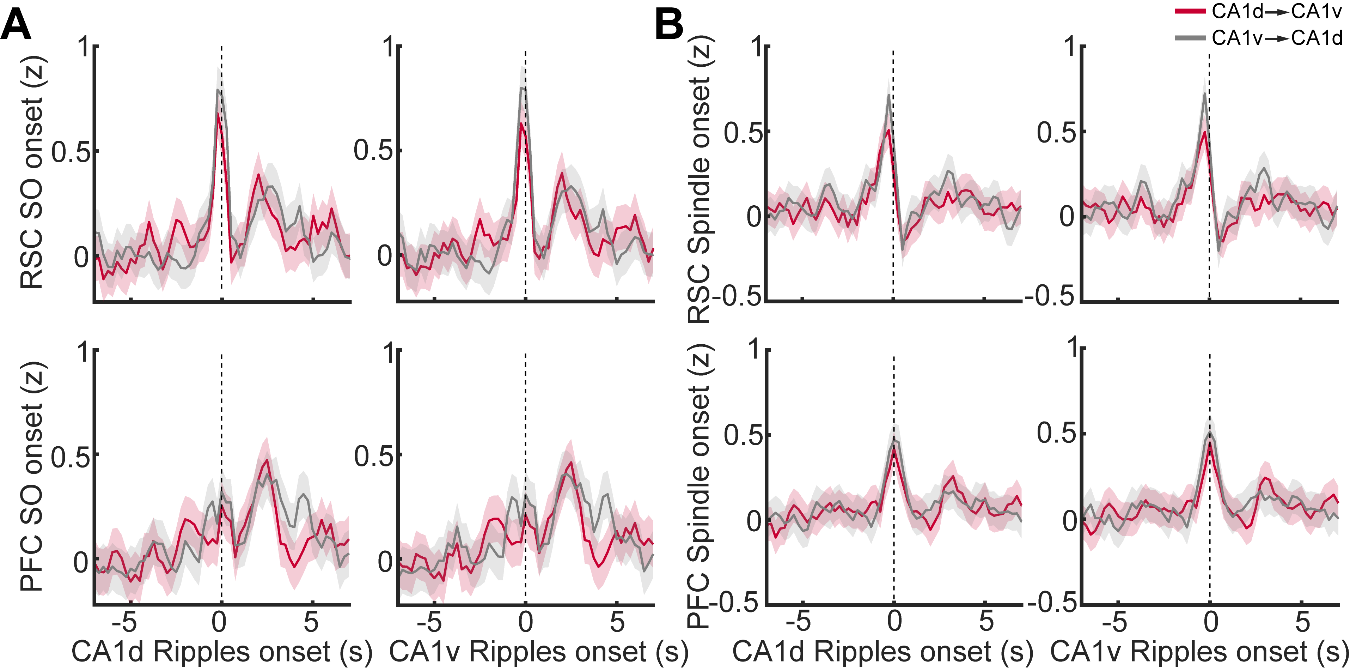

**Figure S6. Modulation of cortical oscillation dynamics by the propagation direction of coordinated hippocampal ripples. A**, average z-scored count of SOs in RSC (top row) and PFC (bottom row) time-locked to the onset (t = 0) of coordinated ripples. Each plot displays SOs aligned to ripple onset in either CA1d (left column) or CA1v (right column), comparing CA1d-to-CA1v (red) and CA1v-to-CA1d (gray) propagation directions. **B**, corresponding plots showing average z-scored power of spindle events in RSC (top row) and PFC (bottom row), time-locked to ripple onset as in (A). Solid lines indicate the smoothed mean, and shaded areas represent the standard error of the mean (S.E.M.). Statistical significance was determined using point-wise two-sided Wilcoxon rank-sum tests with False Discovery Rate (FDR) correction for multiple comparisons. No significant statistical differences were detected (p < FDR-corrected threshold and |Cohen’s d| above criterion) within the analysis window.

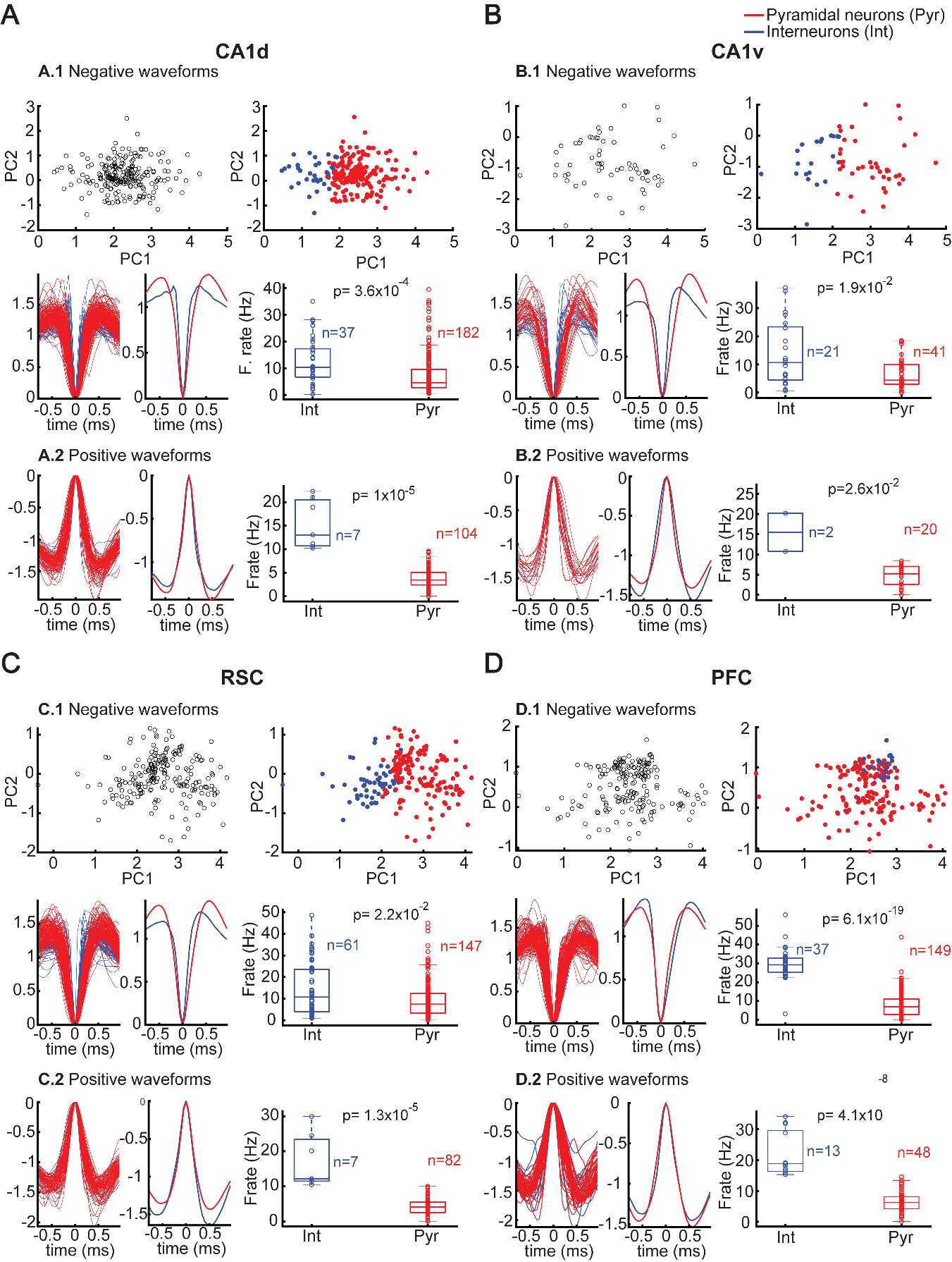

**Figure S7. Classification of single units as putative pyramidal neurons and interneurons across recording sites. (A–D)** Summary of spike sorting and cell-type classification for CA1d (A), CA1v (B), RSC (C), and PFC (D). For each region: (1) Top: Principal component (PC) scatter plots of negative waveform components for all recorded units (left, black), and color-coded separation of putative pyramidal neurons (Pyr, red) and interneurons (Int, blue) based on waveform features (right). (2) Bottom: Overlaid (left) and mean (center) spike waveforms (normalized to minimum amplitude) for each unit type, and boxplots comparing average firing rates between interneurons and pyramidal neurons, with unit counts indicated (n) and p-values from two-sided Wilcoxon rank-sum tests (right). Panels A.2, B.2, C.2, and D.2 display the same waveform and firing rate analyses as the 'bottom plots' for negative waveform units. Note that PC scatter plots are not included for positive waveform units. Classification for positive waveform units was based solely on their firing rate, as clear separation could not be obtained with k-means clustering. In all boxplots, the box indicates the interquartile range (25th–75th percentile), the line marks the median, whiskers depict lower (Q1) and upper (Q4) quartiles, and individual data points are shown as semi-transparent dots.

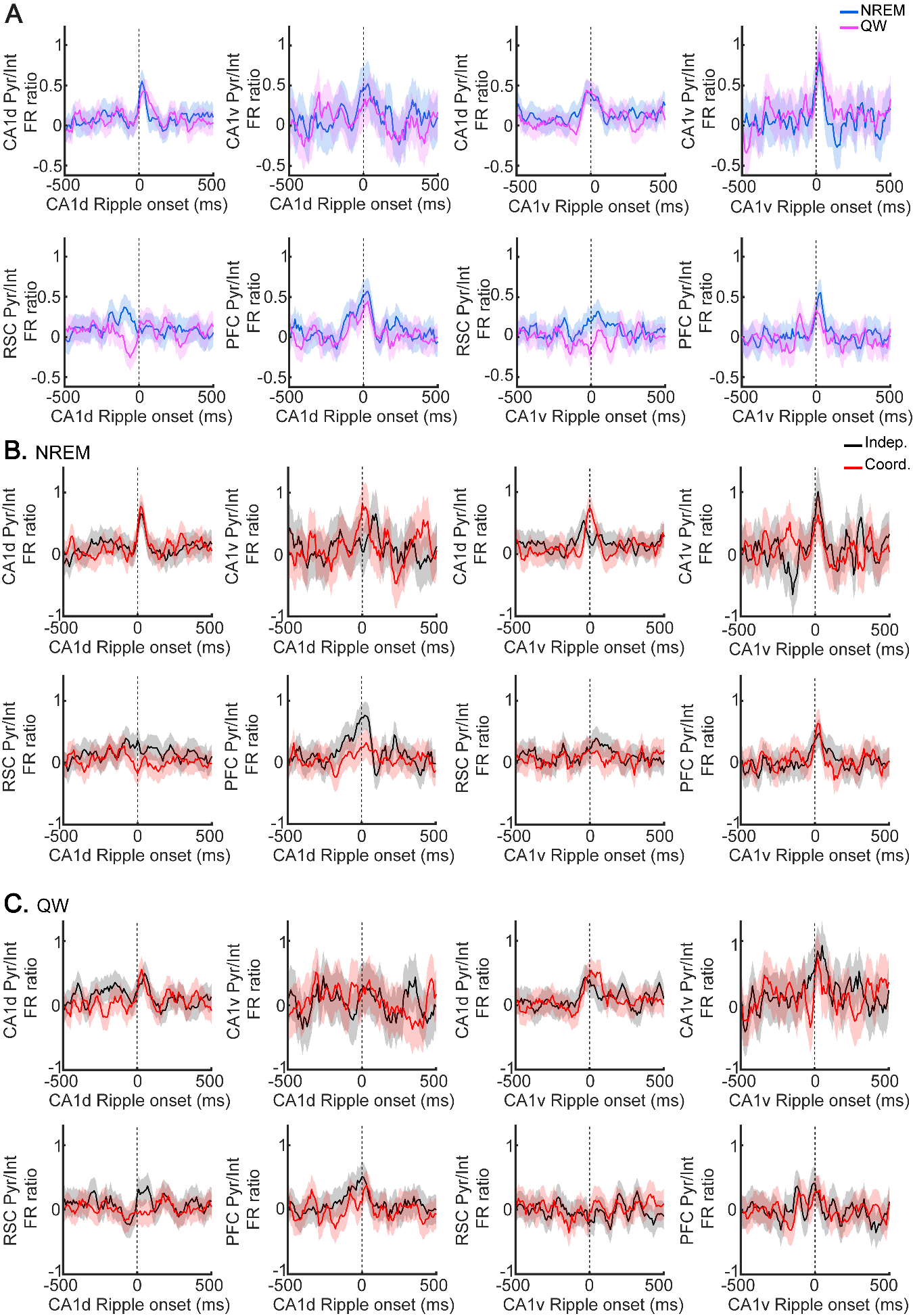

**Figure S8. Dynamics of pyramidal-to-interneuron firing rate ratios aligned to ripple onset across brain regions and brain states.** **A**, time course of the pyramidal/interneuron (Pyr/Int) firing rate ratio in CA1d, CA1v, RSC, and PFC, aligned to ripple onset (t = 0 ms) in CA1d (left columns) and CA1v (right columns), for NREM sleep (blue) and quiet wakefulness (QW, magenta). **B**, same as (A), but during NREM, comparing independent (black) versus coordinated (red) ripples. **C**, same as (B), but during QW. In all panels, solid lines represent the mean ratio, and shaded regions indicate the S.E.M. Statistical significance was determined using point-wise two-sided Wilcoxon rank-sum tests with False Discovery Rate (FDR) correction for multiple comparisons. No significant statistical differences were detected (p < FDR-corrected threshold and |Cohen’s d| above criterion) within the analysis window.

**Supplementary tables**

**Table S1. Histological identification of recorded brain regions per animal.**

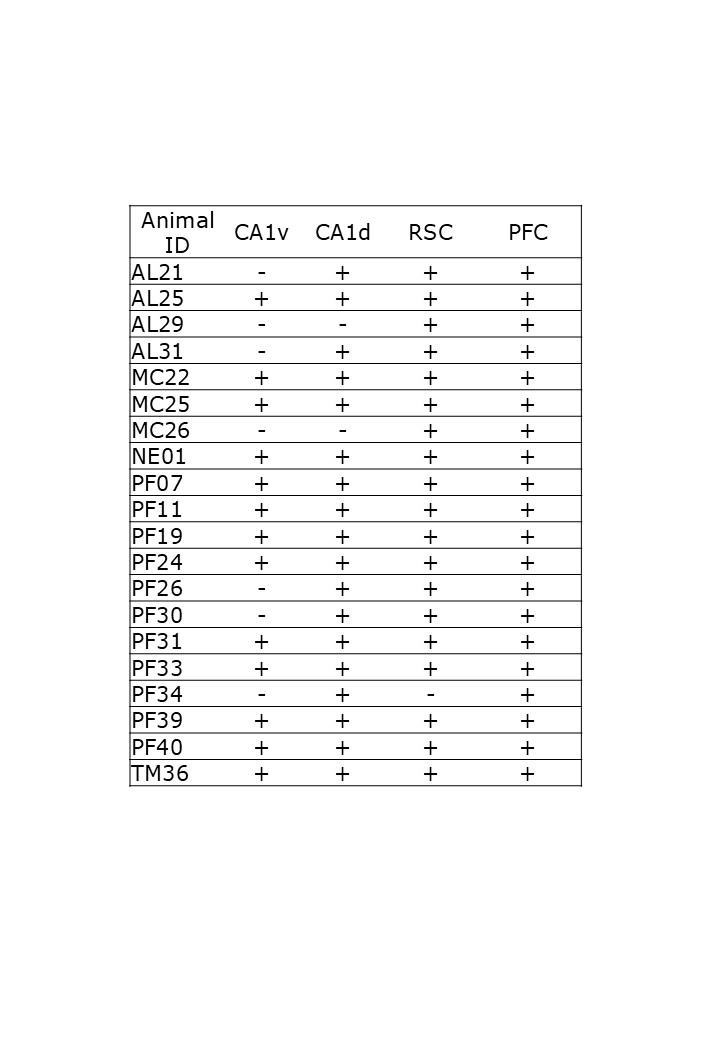

‘+’ indicates successful electrode placement in the region and inclusion of its data in analysis; whereas ‘–’ depicts failed electrode placement.

**Table S2. Duration and relative proportion of brain states across animals during sleep recordings.**

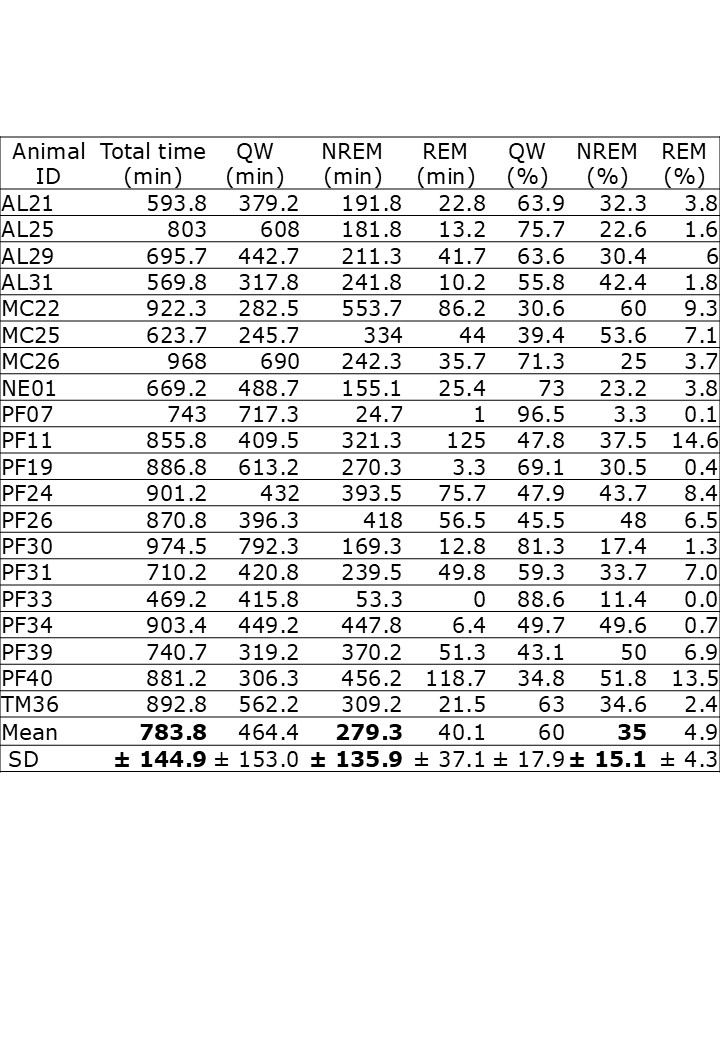

Cumulative recording time (in minutes) per animal, together with the absolute (min) and relative (%) durations of nREM sleep, REM sleep, and quiet wakefulness (QW). Data represents totals across all recorded sessions. nREM mean and relative times are highlighted in bold.

**Table S3. Total recording sessions and unitary activity recorded per animal and brain region.**

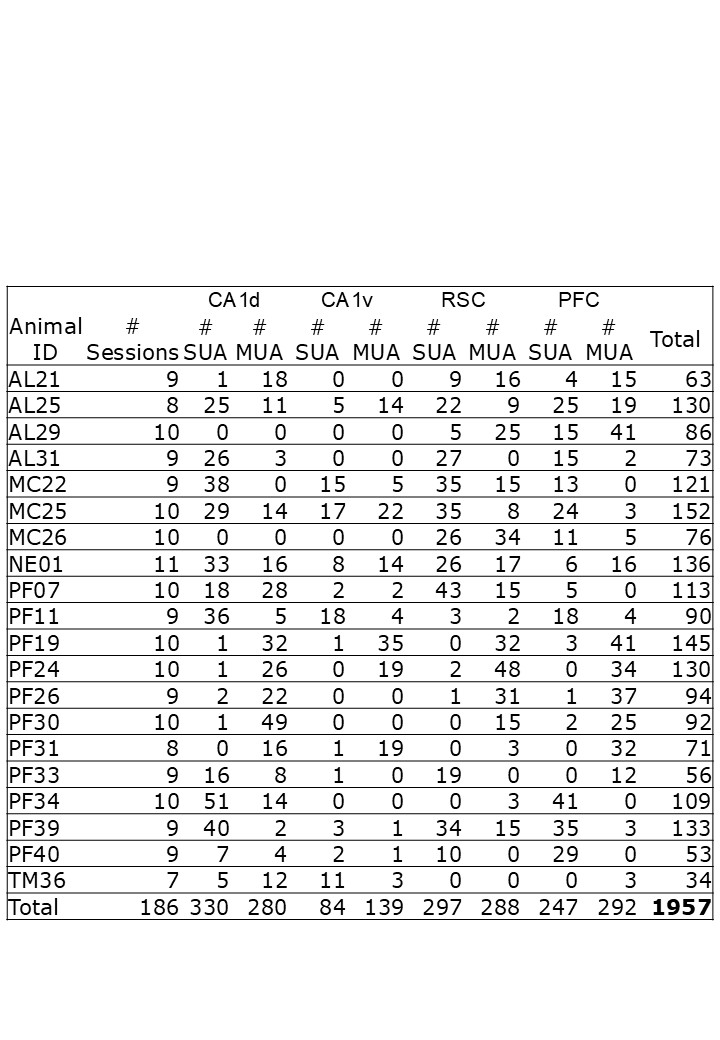

SUA stands for single unit activity. MUA stands for multi-unit activity. Total refers to the sum of all recorded SUAs and MUAs per animal.

**Table S4. Fixed effects coefficients from a linear mixed-effects model of neuronal firing rate during ripple episodes.**

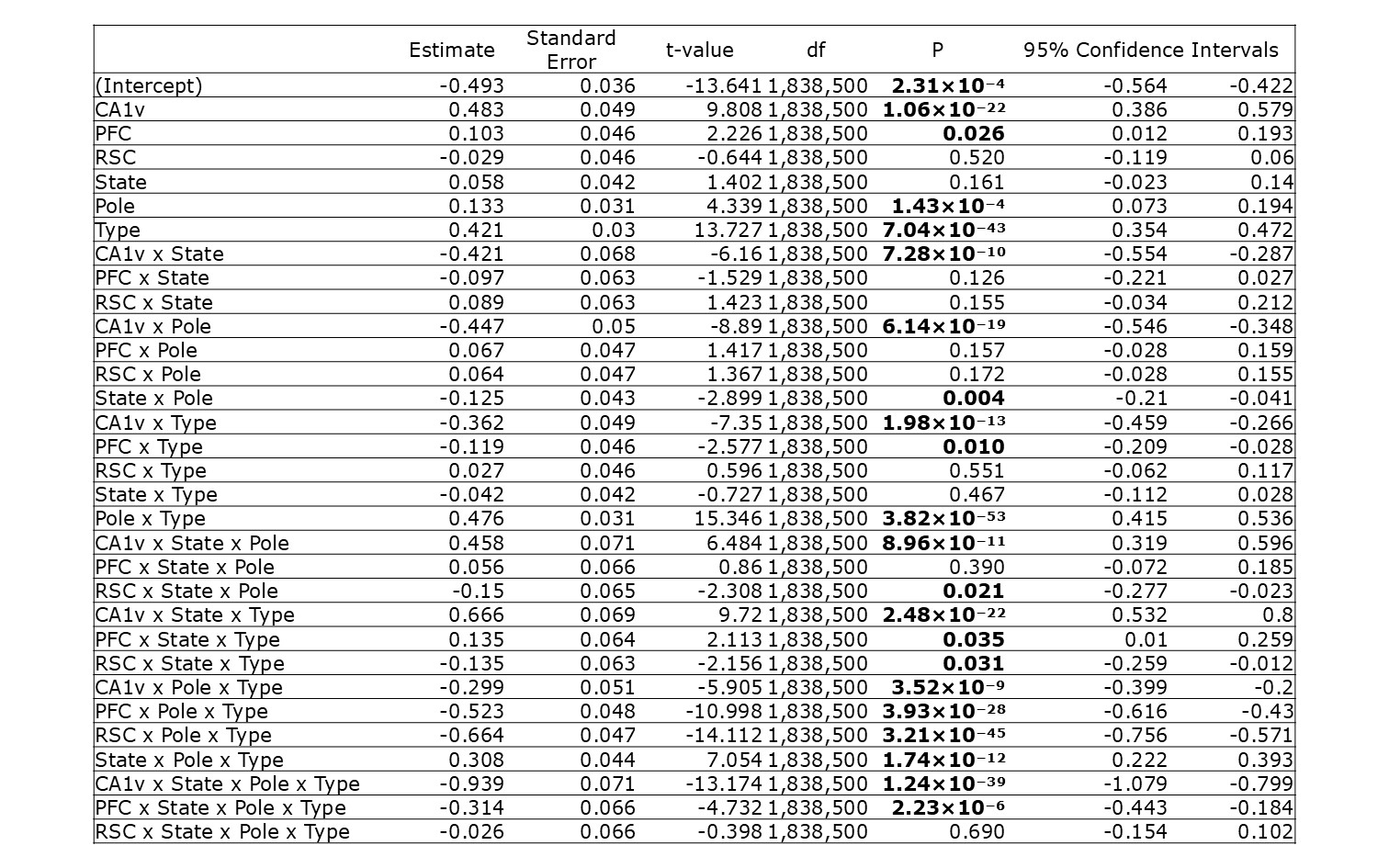

Fixed effects coefficients (Estimate), their standard errors, t-statistics (t-value), degrees of freedom (df), p-values (P), and 95% confidence intervals (confidence limits). Coefficients for CA1v, PFC, and RSC represent the estimated difference in neuronal firing rate compared to the CA1d region, holding other factors constant. The intercept represents the estimated neuronal firing rate when all factors are at their reference levels. State refers to brain state (QW or nREM). Pole refers to ripple initiation pole of the hippocampus (dorsal or ventral). Ripple type refers to ripple synchrony (independent or coordinated). P < 0.05 are highlighted in bold. Firing rate during ripple episodes was modeled as a function of fixed effects (recorded region, brain state, initiation pole, ripple type, and their interactions) and random effects (animal and session), according to the following formula:

Firing rate ~ state * pole * type * region + (1|animal) + (1|animal:session) + residuals

**Table S5. Summary of linear mixed-effects model parameters and fit statistics.**

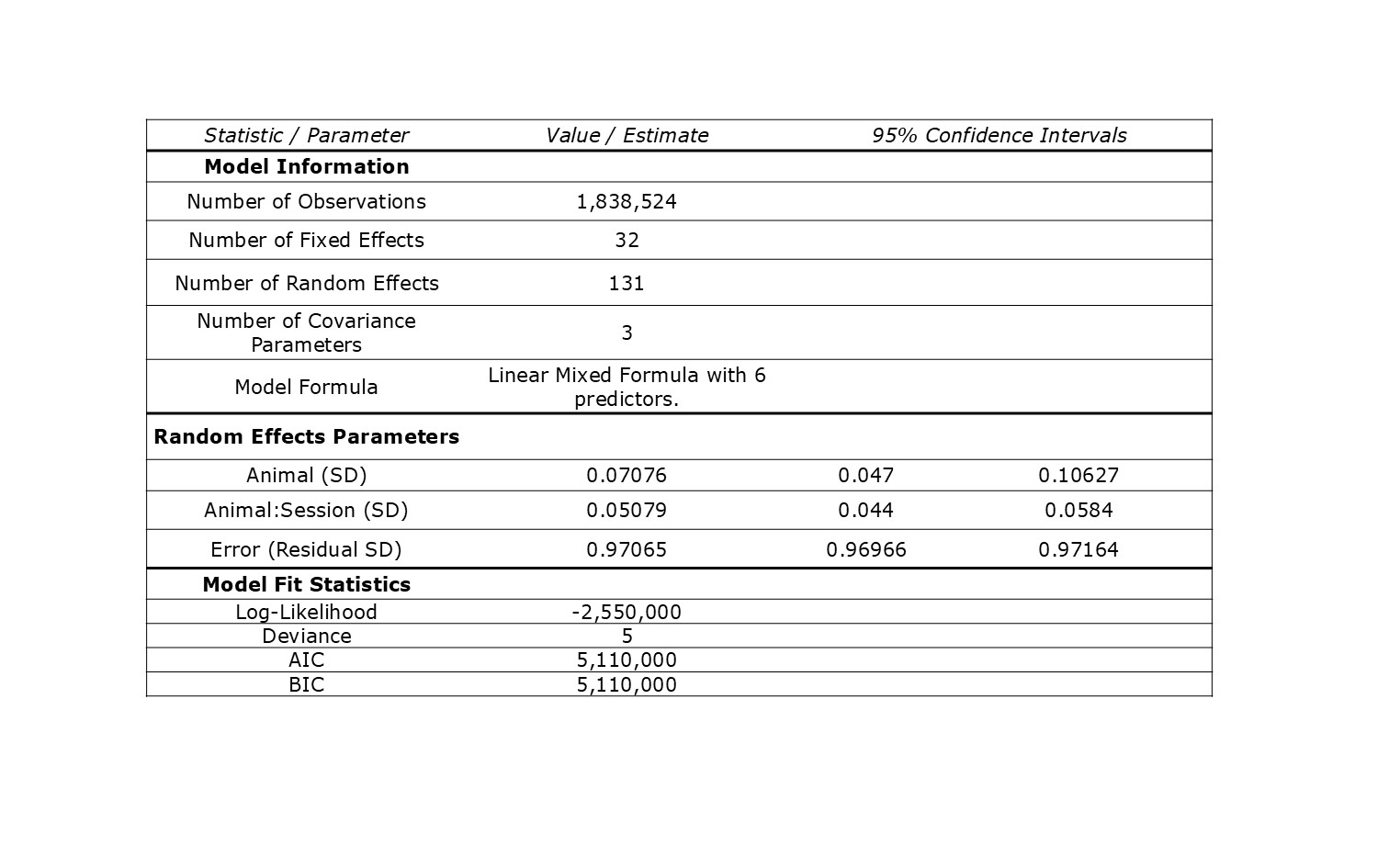

Note: AIC, Akaike Information Criterion; BIC, Bayesian Information Criterion.

**Table S6. Analysis of variance (ANOVA) marginal tests (Type III) for linear mixed-effects model.**

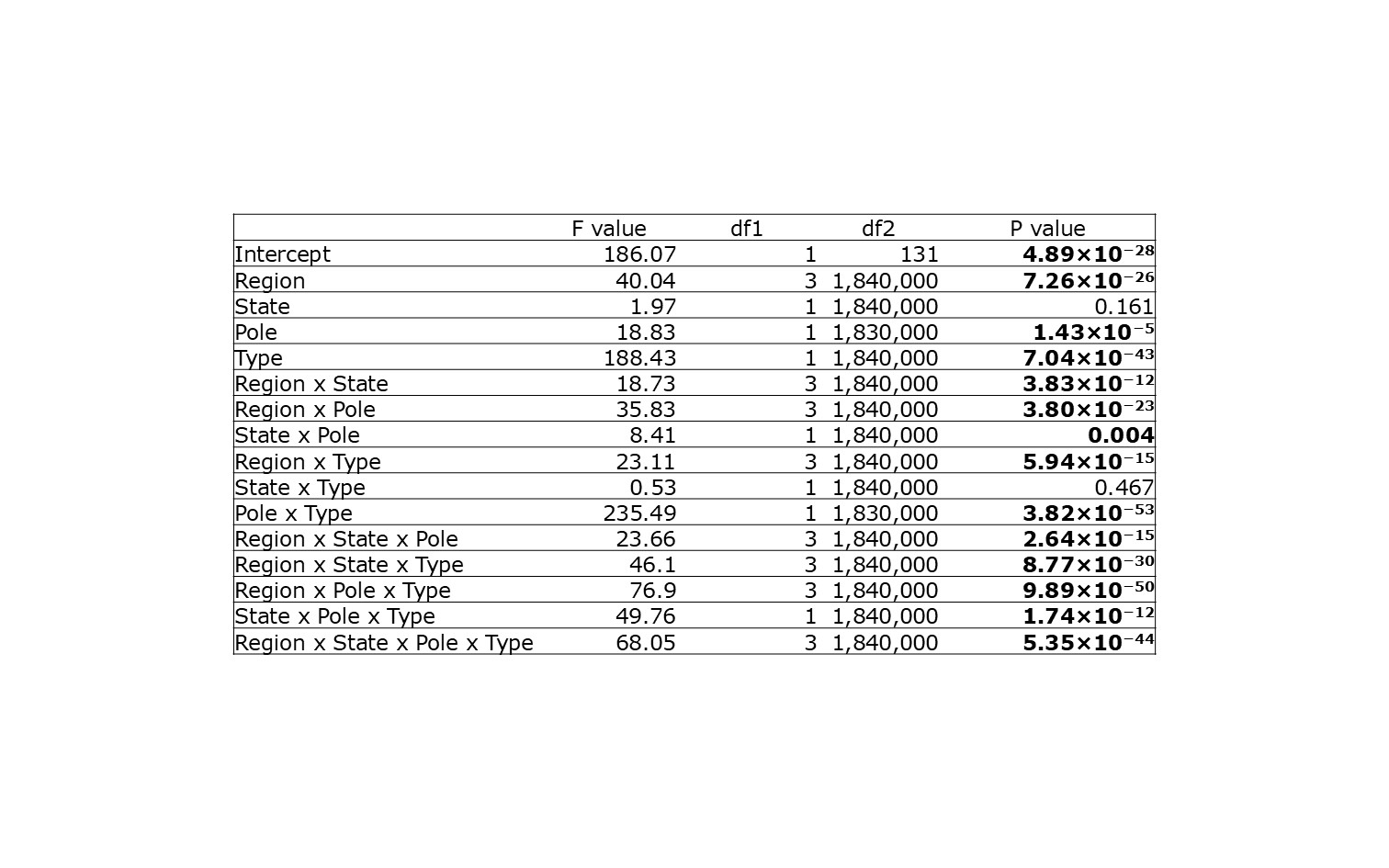

F value represents the variance explained by a factor in relation to the variation expected due to random error. df1 and df2 represent the numerator and denominator degrees of freedom, respectively. The model's marginal R-squared, representing variance explained by fixed effects, was 0.052. The conditional R-squared, representing variance explained by both fixed and random effects, was 0.058. P < 0.05 are highlighted in bold.
